# Extrinsic regulation of interneuron specification and migration

**DOI:** 10.1101/2022.05.03.490384

**Authors:** Fabrizia Pipicelli, Natalia Baumann, Rossella Di Giaimo, Christina Kyrousi, Rebecca Bonrath, Denis Jabaudon, Silvia Cappello

## Abstract

The imbalance between excitatory and inhibitory neurons in the human brain might lead to neurodevelopmental and neuropsychiatric disorders including cortical malformations, epilepsy, and autism spectrum disorders. We propose that the extracellular environment regulates interneuron differentiation and migration during development, ultimately affecting the excitatory/inhibitory balance.

Using ventral cerebral organoids and dorso-ventral cerebral assembloids with mutations in the extracellular matrix gene LGALS3BP, we show that the composition of the extracellular environment regulates the molecular differentiation of neurons, resulting in alterations in migratory dynamics. To investigate how the extracellular environment affects neuronal specification and migration, we characterized the protein content of extracellular vesicles from cerebral organoids carrying a mutation in *LGALS3BP*, previously identified in individuals with cortical malformations and neuropsychiatric disorders. These results revealed differences in protein composition. Interestingly, proteins associated with cell-fate decision, neuronal migration and extracellular matrix composition were altered in mutant extracellular vesicles. Moreover, we show that treatment with extracellular vesicles changes the transcriptomic profile in neural progenitor cells. Our results indicate that neuronal molecular differentiation is regulated by factors released into the extracellular environment.

## Introduction

Neurogenesis and neuronal migration are essential processes for mammalian brain development, especially for the correct assembly of neuronal circuits. Cortical circuitry function relies on adequate excitatory/inhibitory (E/I) balance, coordinated by glutamatergic and GABAergic neuronal activities, such that regulation of migration of these respective cell types during development is critical(Guo and Anton, 2014; Steinecke et al., 2014).

The mammalian neocortex is mainly populated by glutamatergic excitatory neurons, while GABAergic inhibitory neurons (here “interneurons”, INs) are present in smaller proportions(Peyre et al., 2015). In rodents, approximately 15-20% of neocortical neurons are INs(Tatti et al., 2017), while primates, including humans, have higher proportions(Džaja et al., 2014; Krienen et al., 2020).

In humans, excitatory neurons are generated from apical and basal radial glia cells (aRGs and bRGs, respectively) and intermediate progenitors (IPs) located in the germinal zones: the ventricular (VZ), subventricular (SVZ), and outer subventricular zone (OSVZ). Using the basal process of aRGs and bRGs as a scaffold, they migrate radially into the cortical layers. This process relies on the integrity of aRGs, and defects in radial migration can lead to disorganization of cortical layers or ectopic neurons, which are hallmarks of cortical malformations(Fernández et al., 2016; Fietz and Huttner, 2010; Klaus et al.).

While excitatory neurons migrate and remain in the dorsal forebrain, most cortical INs migrate over long distances from their birthplace to their final positions in the neocortex. Specifically, INs are born in the ganglionic eminences (GEs) in the ventral forebrain from the proliferative zone of the medial ganglionic eminence (MGE) and the caudal ganglionic eminence (CGE)(Bajaj et al., 2021; Peyre et al., 2015). As they became post-mitotic, INs migrate tangentially into the neocortex. INs are highly polarized and, during their migration, they extend branches from the leading process. The stabilization of one of the branches leads to cytoskeleton remodeling that results in nucleus displacement in a process called nucleokinesis. This cyclic movement is characteristic of interneuron migrations and, in particular, of their saltatory migratory behavior(Bellion et al., 2005; Peyre et al., 2015; Silva et al., 2018).

The extracellular matrix (ECM), a complex and dynamic cellular microenvironment, plays a crucial role in progenitor proliferation, differentiation, morphogenesis, and neuronal migration during brain development (Long and Huttner, 2019). Specifically, IN migration is guided by extracellular attractive and repulsive cues, such as extrinsic signals, including molecules and proteins secreted in ECM or by cell-cell contact. Eph and ephrins, as well as Robo and Slit and netrins, are attractive and repulsive cues that guide cell migration(Amin and Borrell, 2020; Lim et al., 2018; Marín and Rubenstein, 2001; Peyre et al., 2015).

In this study, we analyze the role of the extracellular environment in IN specification and migration, focusing on the ECM component LGALS3BP. Within the ECM, LGALS3BP interacts with integrins, fibronectins, galectins, laminins, and tetraspanins(Lee et al., 2010; Stampolidis et al., 2015).

We have previously shown that *LGALS3BP* is enriched in human neuronal progenitor cells (NPCs), and its product is secreted via extracellular vesicles (EVs)(Kyrousi et al., 2021). A *de novo* variation in the exon 5 of *LGALS3BP* has been diagnosed in a patient presenting cortical malformations, developmental delay, autism, dysarthria, ataxia, and focal seizures, suggesting its crucial role during brain development. Moreover, we recently showed that LGALS3BP is critical for human corticogenesis, shaping the extracellular environment and regulating NPC delamination and neuronal migration through its extrinsic function(Kyrousi et al., 2021).

Here, using human cerebral organoids and assembloids, we propose that interneuron differentiation is influenced by secreted factors released in the extracellular environment. Our findings suggest that changes in the extracellular environment, given by the *LGALS3BP* variant, are involved in maintaining excitatory/inhibitory (E/I) neurons balance during brain development.

## Results

### *LGALS3BP* E370K-mutant ventral organoids show alteration in cell identity

To investigate if the E/I balance is influenced by secreted factors, we chose LGALS3BP as a model to study this response, mainly because it is secreted and functions during neurodevelopment(Kyrousi et al., 2021).

To this end, we generated ventral forebrain organoids (vCOs) (Fig EV1A and B)(Bagley Joshua A , Reumann Daniel , Bian Shan, 2017). The vCOs serve as a model to investigate cell fate and differentiation of ventral progenitors into INs. The ventral proliferative zones of MGE and CGE generate different subpopulations of INs(Peyre et al., 2015), while LGE mostly gives rise to medium-spiny neurons (MSNs)(Miura et al., 2020) (Fig EV1A). We used an iPSC line carrying the *LGALS3BP* variant in heterozygosity (E370K) previously described(Kyrousi et al., 2021) (Fig EV1C). We, then, generated vCOs from isogenic control and E370K iPSC line. Mutant vCOs are characterized by a significant decreased LGALS3BP expression compared to the control (Fig 1A and B). We then analyzed the expression of typical markers of MGE (NKX2-1), CGE (PAX6), and LGE (MEIS2)(Bagley et al., 2017; Miura et al., 2020; Yu et al., 2021). We observed a significant decrease of NKX2-1+ cells and a significant increase of PAX6+ and MEIS2+ cells in E370K vCOs (Fig 1C and D). This suggests that E370K vCOs generate more CGE and LGE regions; however, PAX6 is also a marker for dorsal cortical progenitors in VZ and SVZ. To understand the regional identity of the PAX6+ cells found in the E370K vCOs, we performed immunohistochemistry for EOMES, dorsal markers of cortical intermediate progenitors (IP), TBR1, marker of deep layer cortical neurons and SATB2, marker of upper layer cortical neurons (Fig 1E). Surprisingly, E370K vCOs express EOMES, TBR1, and SATB2. EOMES labeled cells in 50% of observed ventricles, TBR1 in about 60%, and SATB2 in about 90%, indicating that the E370K vCOs unexpectedly express dorsal cortical markers (Fig 1F). The presence of cortical IP and neurons was combined with a significant decrease in CALB2+ interneurons(Kanton et al., 2019; Yu et al., 2021) in E370K vCOs (Fig 1G and H), showing an imbalance in excitatory/inhibitory neuron proportions.

**Fig 1.**
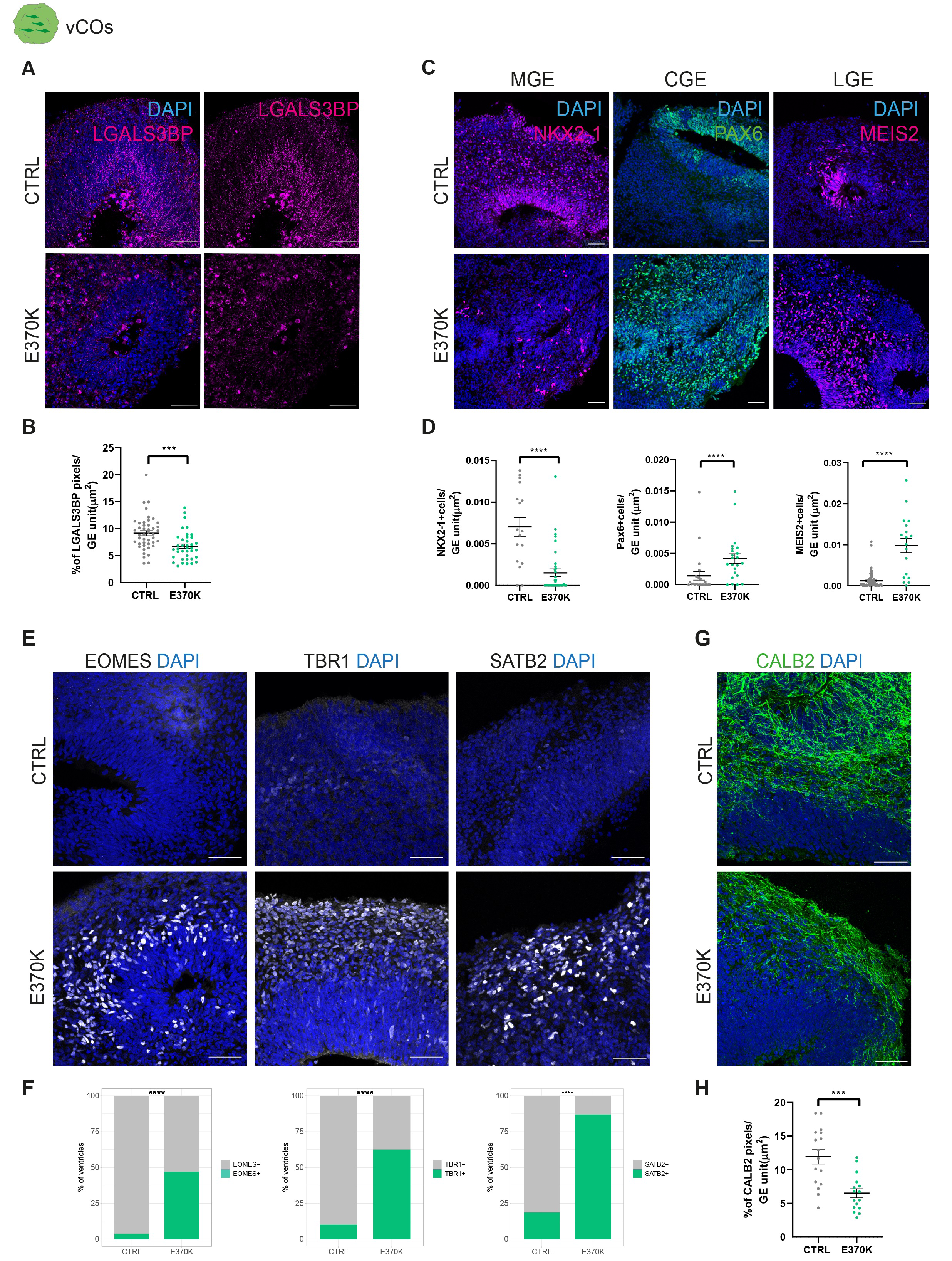
L*G*ALS3BP E370K-mutant ventral organoids show alteration in cell identity A. Representative immunostaining of vCOs for SOX2 (green) and LGALS3BP (magenta). Scale bar: 50 µm. B. Quantification of the percentage of pixel of the LGALS3BP staining in GE unit (µm2). Data are shown mean ± SEM. Statistical significance was based on two-tailed Mann-Whitney U test ****p<0.0001. C. Representative immunostaining of vCOs for MGE marker (NKX2-1, magenta), for CGE marker (PAX6, green) and LGE markers (MEIS2, magenta). Scale bar: 50 µm. D. Quantification of the number NKX2-1, PAX6 and MEIS2 +cells per GE unit (µm2). Data are shown mean ± SEM. Statistical significance was based on two-tailed Mann-Whitney U test ***p<0.001. E. Representative immunostaining of vCOs for cortical markers of IPs (EOMES) of deep layer neurons (TBR1) and upper layer neurons (SATB2). Scale bar: 50 µm. F. Quantification of percentage of ventricles with EOMES+cells (>10 cells) (top), TBR1+cells (>10 cells) (middle) and SATB2+cells (>10cells) (bottom) in the ventral side of vCOs. Statistical significance was based on exact binomial test ****p<0.0001. n of ventricles: for EOMES, CTRL=96, E30K=53; for TBR1, CTRL=100, E30K =107; for SATB2, CTRL=48, E30K =76; from 3 different batches. G. Representative immunostaining of vCOs for CALB2.Scale bar: 50 µm. H. Quantification of the percentage of pixel of the CALB2 staining in GE unit (µm2). Data are shown mean ± SEM. Statistical significance was based on two-tailed Mann-Whitney U test **p<0.01. Every dot in the plots refers to analyzed ventricles per vCO from at least 3 different vCOs generated in at least 2 independent batches.

Moreover, to avoid masking the phenotype due to the functional allele in the heterozygous E370K variant, we also genetically edited an iPSCs line carrying the Y366Lfs variation in *LGALS3BP* in homozygosity(Kyrousi et al., 2021) (Fig EV1C). We then generated Y366Lfs-vCOs. As was the case in E370K vCOs, Y366Lfs-vCOs showed reduced expression of LGALS3BP (Fig EV1D) and expressed EOMES, TBR1, and SATB2, suggesting that functional *LGALS3BP* is essential for the correct specification of INs (Fig EV1E and F). These results show that LGALS3BP plays a role in dorso-ventral patterning, regulating cell differentiation in the ventral forebrain.

### *LGALS3BP* variation causes alteration in cell fate and developmental trajectory

To dissect the transcriptional signatures of altered cells in the E370K-vCOs, we performed single-cell RNA-sequencing (scRNA-seq) analysis in 60-day old CTRL-vCOs and E370K-vCOs. The 5369 identified cells clustered into eight main groups (Fig 2A and B), including progenitors expressing *TOP2A* and IP expressing *NKX2-1, ASCL1* and *PAX6*, and neurons expressing *MAP2* and *DLX5* (Fig 2C). We also identified cortical genes, such as *TBR1, NEUROG1, GLI3* and *SLC17A6* being expressed in E370K vCOs (Fig 2C and Fig EV2A), confirming their dorsal identity also at the molecular level.

**Fig 2.**
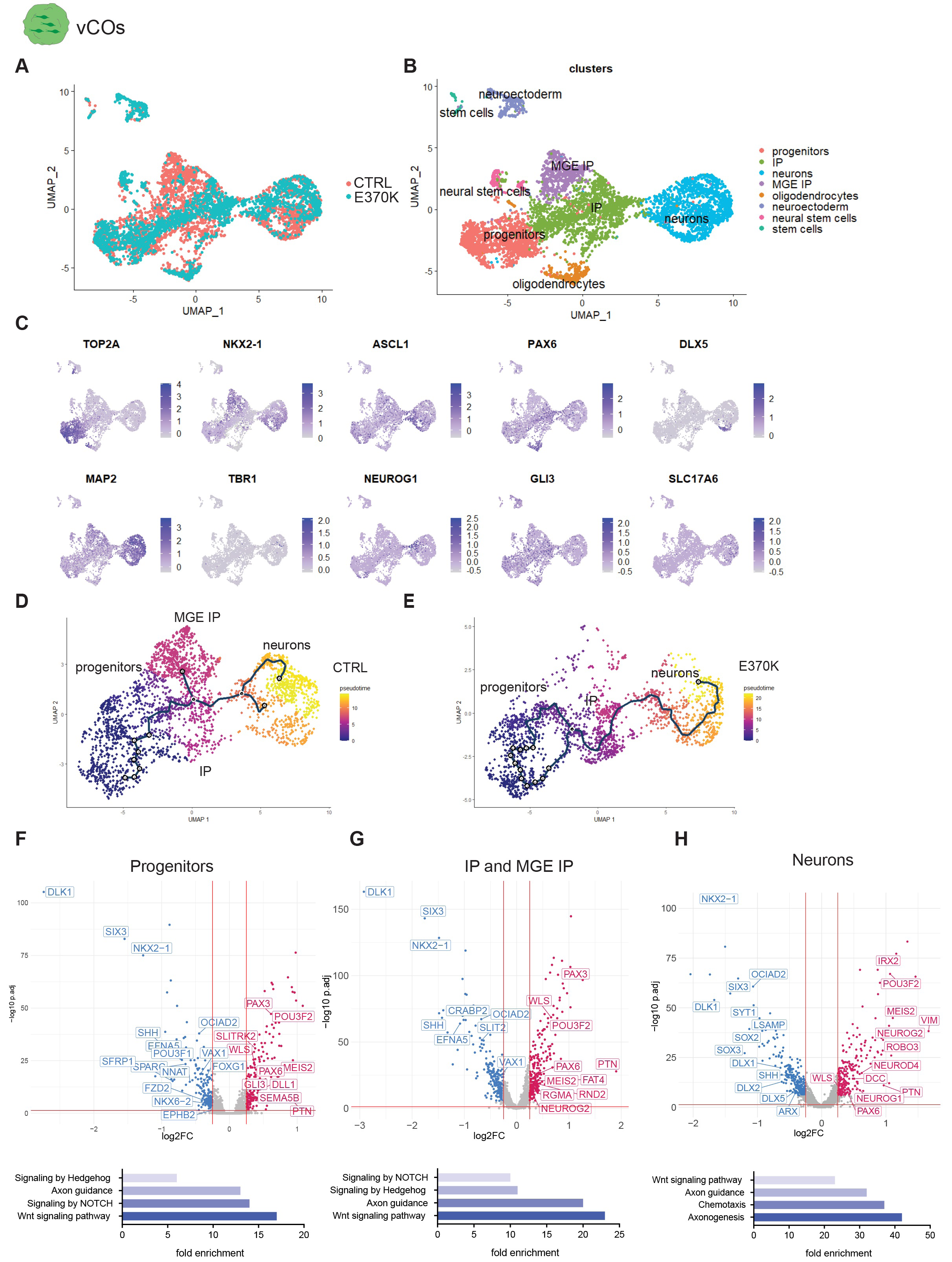
L*G*ALS3BP variation causes alteration in cell fate and developmental trajectory A-B UMAP visualization of scRNA-seq clusters of control and E370K vCOs (n= 4898 cells from a pool of 5 organoids each condition). C. Feature plot depicting the expression of progenitor markers (TOP2A), IPs markers (TTYH1, NKX2- 1, ASCL1, PAX6) and neurons (DLX5, MAP2, TBR1, NEUROG1, GLI3, SLC17A6) in vCOs. D-E. UMAP visualization of pseudo-time trajectories in control vCOs and in E370K from progenitors to INs. F-H Volcano plot showing the fold change (CTRL vs E370K) of gene expression (top) and GO terms of dysregulated pathways (bottom) in progenitors, IP and MGE IP, and in neurons in vCOs.

To investigate if the LGALS3BP variation could cause developmental trajectory alteration, ventral telencephalic cells were aligned on a developmental pseudo-differentiation showing the trajectory from progenitors to neurons (Fig EV1B and C). Using Monocle3(Trapnell et al., 2014), (https://cole-trapnell-lab.github.io/monocle3/), we identified pseudotime trajectories in both CTRL-vCOs and E370K-vCOs (Fig 2D and E and Fig EV2D and E). As expected from the decrease of NXK2-1+ MGE IP (Fig 1C and D and Fig EV2F-H) in E370K-vCOs, the IP-MGE IP trajectory is missing in mutant organoids.

Next, we performed differential expression (DE) analysis in progenitors, IP and MGE-IP, and neurons. Interestingly, all the three populations of E370K-vCOs downregulated transcription factors (TFs) - such as *NKX2-1*, *NKX6-2*, *SIX3*, *OCIAD2 -* and the secreted molecule *SHH,* associated with a ventral patterning. On the other hand, E370K-vCOs upregulate dorsal TFs, like, *PAX6*, *PTN*, *POU3F2*, *GLI3,* and *NEUROG2,* and the secreted molecule *WLS* confirms the dorsal identity of E370K ventral cells. In particular, we found dysregulation of the WNT, NOTCH, and HEDGEHOG pathways, known to regulate cell differentiation and dorso-ventral patterning(Hayward et al., 2008). We also identified molecules associated with axon guidance: *EFNA5*, *EPHB2*, *SEMA5B,* and *SLIT2.* Interestingly, also *FAT4*, *DLL1,* and *RND2*, genes previously associated with periventricular heterotopia (PH)(Klaus et al.), were dysregulated in E370K-vCOs (Fig 2 H-J), suggesting that the cell fate switch can affect the migration of neurons. Mutant neurons show downregulation of interneuron markers, such as *DLX1, DLX2, DLX5*, while they upregulate cortical markers like *NEUROG1, NEUROG2, NEUROD4*, (Fig 2H) confirming the excitatory/inhibitory unbalance found in E370K vCOs.

The DE analysis revealed that E370K-vCOs dysregulate genes are involved in pattern specification, regionalization, differentiation, and neurogenesis as indicated from the enriched GO terms (Fig. EV 2I- K), suggesting a specific contribution of LGALS3BP in human ventral forebrain development. The alteration in expression of secreted and transmembrane molecules, such as SHH, WLS, EPHB2, and SLIT2, in E370K vCOs, show that cell fate might be regulated in an extrinsic way.

### Cell fate changes result in migratory defects

Excitatory and inhibitory neurons migrate in different ways(Buchsbaum and Cappello, 2019; Silva et al., 2018), and, an altered cell fate could lead to migratory defects.

To investigate how E370K vCOs with dorsal identity migrate, we generated human dorso-ventral cerebral assembloids (dvCAs), a suitable human model system to study interneuron migration by resembling the ventral-dorsal forebrain axis in vitro(Bagley Joshua A , Reumann Daniel , Bian Shan, 2017; Birey et al., 2017; Xiang et al., 2017) (Fig EV1A). dvCAs express typical ventral forebrain markers such as NKX2-1 in the ventral region (vCAs) and dorsal forebrain markers such as TBR1 in the dorsal region (dCAs) (Fig EV1B).

We then generated dvCAs from isogenic control and genetically edited iPSCs, *LGALS3BP*-E370K (E370K) (Fig 3A). We generated control dvCAs (vCTRL-dCTRL), and dvCAs with mutant ventral side (vE370K-dCTRL) to focus on the migration of cells with the *LGALS3BP* variation (Fig 3A).

**Fig 3.**
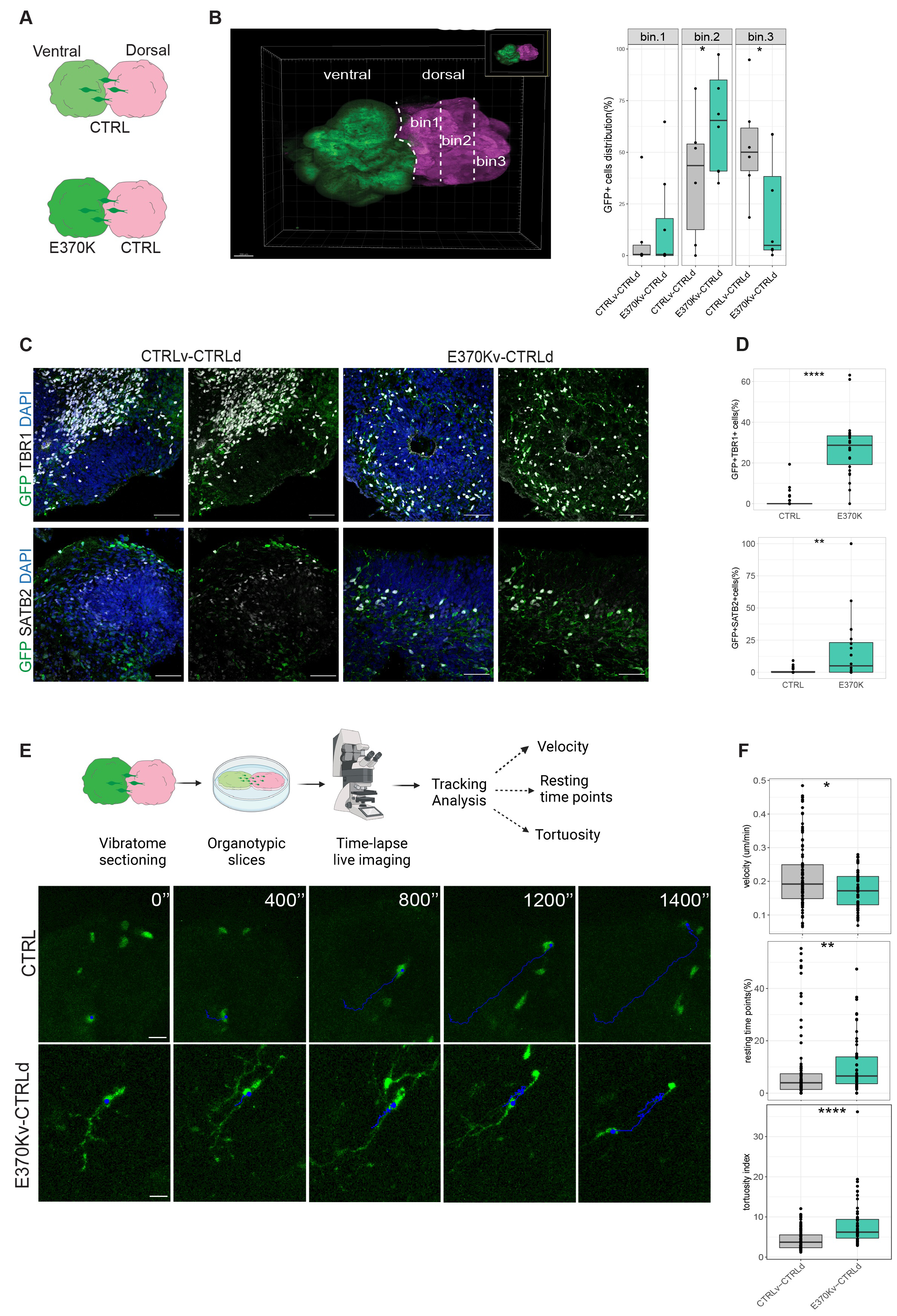
Cell fate changes result in migratory defects A. Schematic of experimental set-up showing different combinations of dvCAs. B. Binning analysis in cleared dvCAs. Scale bar: 200 µm (left) and quantification of distribution of GFP+ventral cells migrated from ventral to dorsal in dvCAs (right). Box plots show median and interquartile range. Statistical significance was based on the Mann-Withney U test *p<0.05. Every dot in the plots refers to analyzed vCOs generated in at least 2 independent batches. C. Representative immunostaining of dorsal regions of dvCAs showing ventral migrated cells (GFP) and neurons (TBR1 and SATB2, white). Scale bar: 50 µm. D. Quantification of the percentage of migrated GFP ventral cells expressing TBR1 and SATB2 in dvCAs. Box plots show median and interquartile range. Significance was based on the Mann-Withney U test **p<0.01, ****p<0.0001. Every dot in the plots refers to analyzed ventricles per vCO from at least 3 different vCOs generated in at least 2 independent batches. E. Schematic overview of experimental set-up for time-lapse live imaging in dvCAs (top) and examples of the GFP+ ventral cells movements in dorsal region of dvCAs, monitored for 48h (bottom). Scale bar: 80 µm. F. Quantification of velocity (top), number of resting time points (middle) and tortuosity index (bottom) of migrating GFP+ventral cells. Box plots show median and interquartile range. Statistical significance was based on the Mann-Withney U test *p<0.05, **p<0.01, ****p<0. 0001.Every dot in the plots refers to single cells per vCO.

To monitor migrating neurons, we first analyzed neurons migrated into the dorsal region, by quantifying the number of cells migrating from the ventral (GFP+, in green) to the dorsal (RFP+, in magenta) region in dvCAs (Fig 3B). Despite an absence of significant differences in the total number of migrated cells (Fig EV3B), we observed significant changes in their distribution within the dorsal region of CAs. We analyzed the distribution of GFP+ cells migrated from the ventral to the dorsal region of CAs by subdividing the dorsal region (dCAs) into three equally distributed regions (bins in Fig 3B). In control dvCAs, GFP+ migrating cells mainly were distributed among bin2 (∼45%) and bin3 (∼55%) of the CTRL-dCAs. In the vE370K-dCTRL CAs, most of the cells (70%) were found in bin2 and only 10% in bin3 of CTRL-dCAs, showing a different distribution of mutant cells when they migrate into the dorsal region, probably because of their altered molecular identity. We then quantified the percentage of ventral E370K GFP+ cells that expressed TBR1 or SATB2 in CTRL-dCAs (Fig 3C) to assess if ventral E370K neurons expressing cortical markers migrate to dCAs. Interestingly, 30% of total E370K GFP+ migrated cells express TBR1 while 10% express SATB2. On the contrary, CTRL GFP+ ventral cells were absent in CTRL-dCAs, as expected (Fig 3D). We additionally quantified TBR1+ cells in CTRL-vCAs and E370K-vCAs and the TBR1+GFP+ cells in CTRL-dCAs after clearing(Masselink et al., 2019) (Fig EV3B). We confirmed the presence of both TBR1+ cells in E370K-vCAs (Fig EV3C) and GFP+TBR1+ cells migrated from E370K-vCAs in the CTRL-dCAs (Fig EV5D).

We showed that scRNA-seq data from mutant neurons revealed alteration of secreted molecules associated with axon guidance (Fig 2H), suggesting changes in migratory dynamics. To dissect the migratory dynamics and behavior of E370K neurons, we monitored the trajectories of GFP+ cells in dCAs. We performed time-lapse imaging of 60-day old dv-CAs slices and tracked the GFP+ ventral cells during their migration within dCAs (Fig 3E). We measured velocity (speed of migration), resting timepoints (time cells spend without moving), and tortuosity (the ability to move in a straight trajectory) as previously described in Klaus et al., 2019(Klaus et al.) (Fig 3F). For all three parameters, we observed a significant difference compared with the control. We found that speed of migration decreased, and resting timepoints and tortuosity increased, as previously shown in the case of mutations of the PH genes *DCHS1* and *FAT4*(Klaus et al.). Interestingly, not only neuronal dynamics but also similar genes (*ROBO3*, *GNG5*, *DCC)* were altered in E370K ventral neurons and DCHS1 and FAT4 altered neurons(Klaus et al.), suggesting common signatures for neurons associated with PH (Fig EV3D and E). In support of their fate switch, the average speed of the E370K ventrally-generated neurons is similar to the one of previously observed dorsally-generated control neurons analyzed in Klaus et al., 2019(Klaus et al.) (Fig EV3F). Altogether, these results show that cell fate changes, observed in the E370K cells, lead to migratory dynamics alteration, especially in directionality, sliding movement, and speed.

### Extrinsic effect of LGALS3BP in ventral progenitor fate and neuronal specification

In previous work, we showed that the addition of control culture media to E370K-COs rescues the phenotype observed, restoring the number of proliferating cells, the thickness of the apical belt, and the localization of neurons(Kyrousi et al., 2021). This finding highlights the extracellular function of LGALS3BP, supporting the important role of the extracellular environment in brain development, suggesting that secreted molecules and proteins regulate cell proliferation, delamination, and distribution.

To assess if the dorsal identity acquired by the mutant cells can be reverted by providing a more physiological extracellular environment, we generated ventral mosaic organoids (vMOs), containing both isogenic control iPSCs and GFP+ E370K IPCs (Fig 4A).

**Fig 4.**
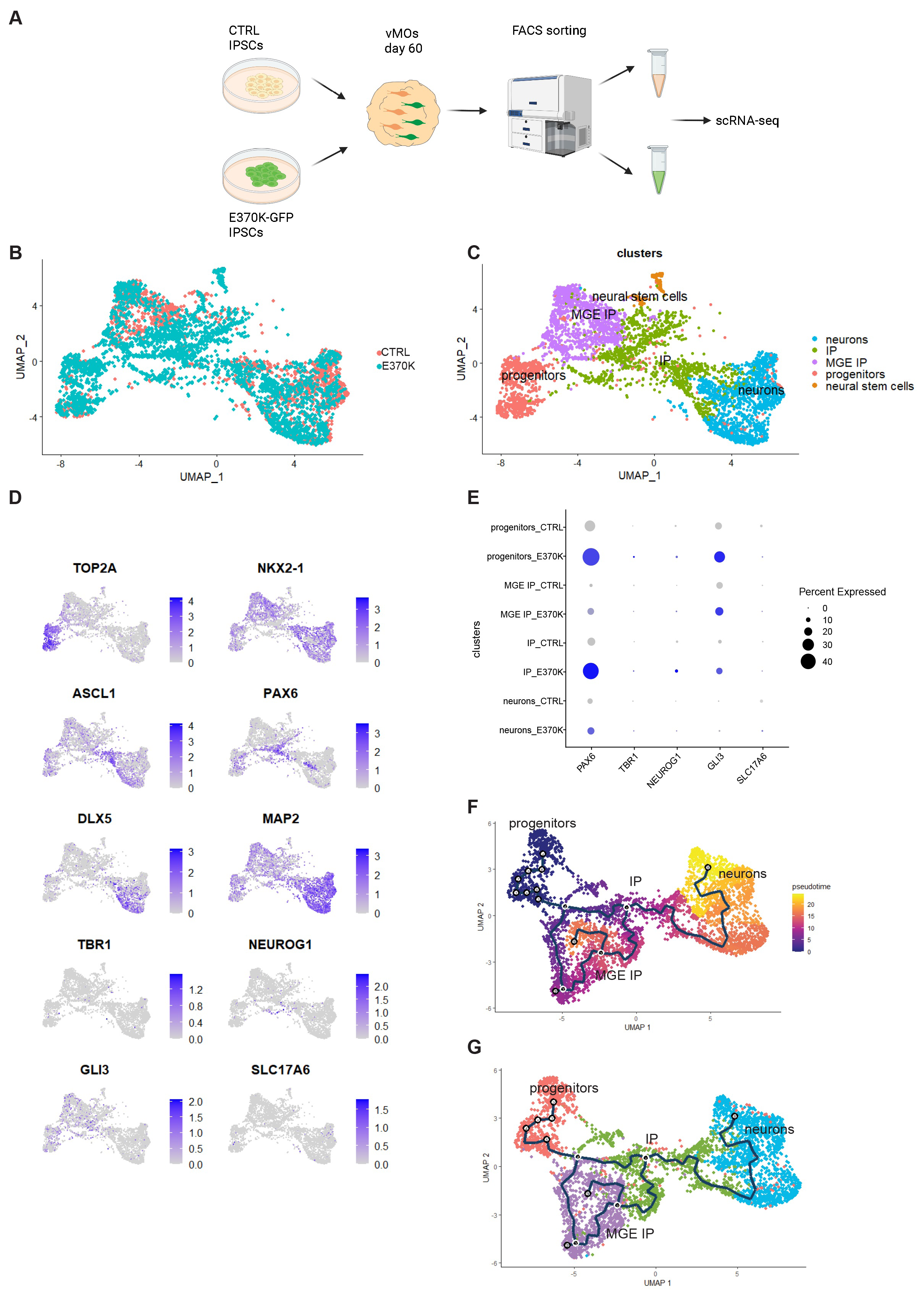
Extrinsic effect of LGALS3BP in ventral progenitor fate and neuronal specification A. Schematic of experimental set-up of generation and processing of vMOs. B-C. UMAP visualization of scRNA-seq clusters of vMOs (n= 5369 cells from a pool of 5 organoids). D. Feature plot depicting the expression of progenitor markers (TOP2A), IPs markers (TTYH1, NKX2- 1, ASCL1), neurons (DLX5, MAP2, TBR1, NEUROG1, GLI3, SLC12A6) in vMOs. E. Dotplot of cortical genes expressed in vMO clusters. F-G. UMAP visualization of pseudo-time trajectories in vMOs from progenitors to INs.

We performed scRNA-seq analysis on vMOs, and the 4898 identified cells were clustered into five main clusters (Fig 3B and C), including progenitors expressing *TOP2A*, IP expressing *NKX2-1 ASCL1*, *PAX6*, neurons expressing *DLX5* and *MAP2*(Bajaj et al., 2021; Yu et al., 2021) (Fig 4D). Cortical genes such as *TBR1, NEUROG1, GLI3* and *SLC17A6*- are also expressed in vMOs, but at lower levels compared to E370K vCOs (Fig 4D and E and Fig 2C and Fig EV2A). As for the E370K vCOs, E370K progenitors, IP and MGE-IP obtained from vMOs express higher level of dorsal genes such as *PAX6* and *GLI3* compared to the control. Interestingly, gene expression of cortical markers such as *TBR1*, *NEUROG1* and *SLC17A6* is decreased in E370K neurons generated from vMOs, showing a partial rescue of the neuronal identity affected in the mutant vCOs (Fig 4D and E, Fig 2C and Fig EV2A).

The identified ventral telencephalic cells were aligned on a developmental pseudotime showing the trajectory from progenitors to neurons, as seen in vCOs (Fig EV4B and C). Using Monocle3(Trapnell et al., 2014), (https://cole-trapnell-lab.github.io/monocle3/), we identified pseudo-time trajectories in both CTRL and E370K cells obtained from the vMOs (Fig 4F and G). The pseudotime trajectories were populated by both CTRL and E370K cells and resemble the ones found in CTRL-vCOs, suggesting a partial rescue of the E370K cells exposure (due to the close proximity) to CTRL cells in the vMOs.

Specifically, the cell proportions of progenitors, IP, and MGE IP are increased compared to the E370K- vCOs (Fig EV4D-F and Fig EV2F-H). In E370K-vMOs we observed an increase of 30% of cells in progenitors, 25% in IP, and 50% in MGE IP compared to the E370K-vCOs, indicating a change also in cell proportions when cells differentiate in a more physiological environment.

These results suggest an extrinsic regulation of neuronal cell fate and differentiation. The ventral molecular identity rescue observed in the E370K cells was guided by the presence of CTRL neighboring cells. On the contrary, both progenitors and IP retained their dorsal identity, downregulating *NKX2-1* and *SHH*, suggesting an extrinsic control over neuronal specification. This is in line with the progressive specification hypothesis, which states that interneurons are restricted to a particular subtype at birth, but their definitive identities are established later in development, and they are shaped by the extracellular (cortical) environment(Kepecs and Fishell, 2014; Wamsley and Fishell, 2017).

### LGALS3BP can revert the molecular identity of mutant ventral progenitors and interneurons

To investigate the molecular identity of E370K vCOs and vMOs, we built a dorso-ventral model that predicts the dorsal or ventral molecular identity of cells based on their transcriptomic features, defining the dorso-ventral score (DV) (Fig EV5A). We performed cluster analysis of scRNA-seq transcriptome data from CTRL-dCOs and CTRL-vCOs, E370K-vCOs, and CTRL/E370K-vMOs (Fig 5A) and identified three main clusters: progenitors expressing *TOP2A*, IP expressing *TTYH1*, and neurons expressing *MAP2* (Fig 5B-D and Fig EV5B).

**Fig 5.**
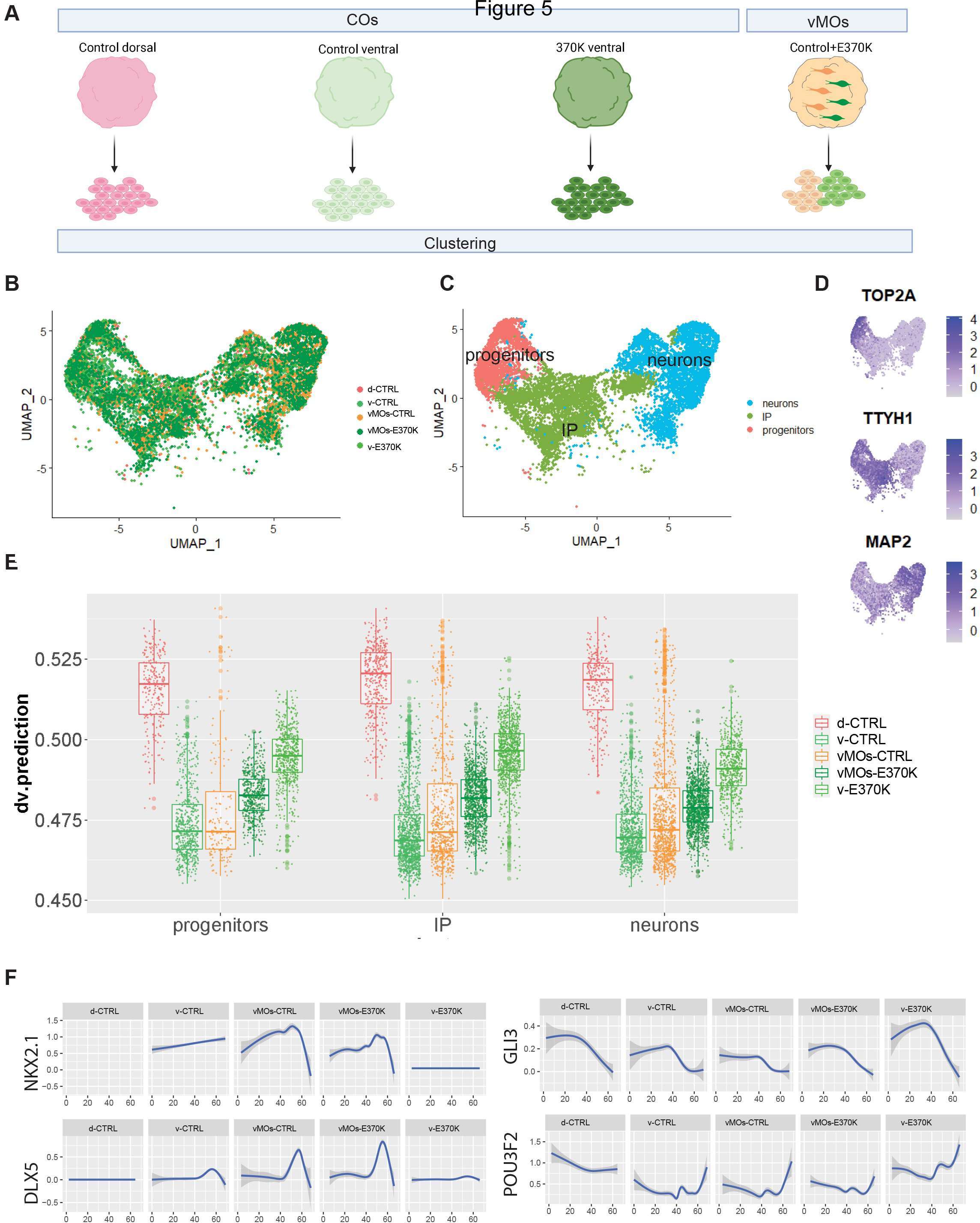
LGALS3BP can revert the molecular identity of mutant ventral progenitors and neurons A. Schematic of experimental set-up of cluster analysis of CTRL dCOs (d-CTRL), CTRL vCOs (v- CTRL), E370K vCOs (v-E370K) and vMOs (vMOs-CTRL and -E370K). B-C. UMAP visualization of scRNA-seq clusters of CTRL dCOs, CTRL vCOs, E370K vCOs and vMOs. D. Feature plot depicting the expression of progenitor marker (TOP2A), IPs marker (TTYH1,), neuronal marker (MAP2). E. Boxplot showing the dorso-ventral prediction score of progenitors, IP, and neurons in all conditions. Box plots show median and interquartile range. F. Expression level of ventral (NKX2-1 and DLX5) and dorsal (GLI3 and POU3F2) genes along the pseudo-time axis in each condition.

As expected, the dorso-ventral prediction model revealed the highest DV score in CTRL-dCOs and the lowest in control CTRL-vCOs (Fig 5E). All the other conditions showed blended dorso-ventral identity and were distributed according to extrinsic short-distance exposure. E370K-vCOs cells mapped midway through the dorso-ventral trajectory while cells from the vMOs were mapping similarly. As expected, CTRL cells from vMOs were closer to CTRL cells from vCOs, while E370K cells from vMOs were closer to E370K-vCOs. These data strongly indicate that E370K cells acquire a dorsal molecular identity, and this can be reverted by exposure to CTRL cells. Finally, CTRL cells from vMOs show mostly a ventral score with some outlier cells with a higher DV score, suggesting that also CTRL cells have been affected by the extracellular environment generated by the neighboring E370K cells (Fig 5E and Fig EV5C and D).

Gene expression is affected in both CTRL and E370K cells from vMOs. *NKX2-1* and *DLX5,* which are almost no detectable or have low expression levels in E370K-vCOs, are expressed along the pseudo-differentiation axis in E370K cells from vMOs (Fig 5F). *GLI3* and *POU3F2*, which are mostly expressed in CTRL-dCOs, are highly expressed in E370K-vCOs, but lower in E370K cells vMOs, without showing great changes in CTRL cells from vMOs (Fig 5G).

To map CO single-cell transcriptome data to the developing mouse brain, we then used VoxHunt(Fleck et al., 2021) (https://quadbiolab.github.io/VoxHunt/) obtained from situ hybridization data (Allen Developing Mouse Brain Atlas, Thompson et al., 2014). As suspected, transcriptome data from CTRL dCOs and vCOs, map, respectively, to the mouse cortex (pallium) and ventral forebrain (subpallium). Data from E370K-vCOs, map to the ventral forebrain but also to the mouse cortex, confirming their dorsal molecular identity. CTRL and E370K cells from vMOs, mostly map to the ventral forebrain; however, E370K cells still show a correlation with the mouse pallium (Fig EV5E). Altogether, these findings show that the molecular identity of progenitors, intermediate progenitors, and neurons is influenced by extrinsic factors.

### Extrinsic function of LGALS3BP in progenitor specification and neuronal migration

During brain development, neurogenesis and cell migration are influenced by extrinsic factors released in the ECM, also via EVs(Peruzzotti-Jametti et al., 2021; Sharma et al., 2019; Taverna and Huttner, 2010). Our previous work indicated that LGALS3BP is secreted in EVs in human cerebral organoids and has a crucial role in modulating the extracellular space(Kyrousi et al., 2021). To investigate the extracellular mechanisms underlying cell fate switch and neuronal migration found in E370K dvCOs, we performed proteomic analysis of EVs collected from CTRL and E370K vCOs (Fig 6A and Fig EV6A).

**Fig 6.**
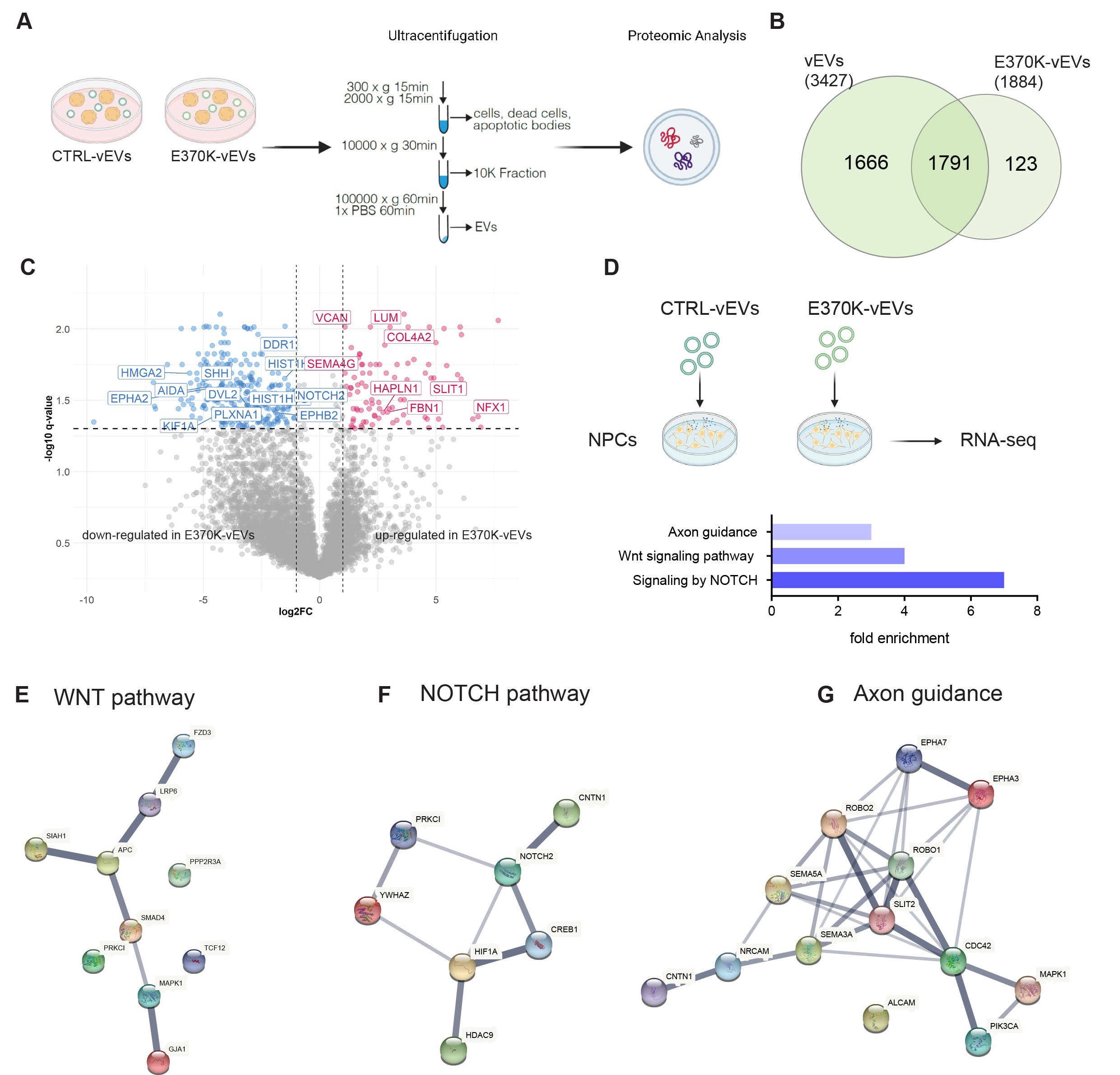
Extrinsic function of LGALS3BP in progenitor specification and neuronal migration A. Schematic of experimental set-up of proteomic analysis of EVs collected from vCOs. B. Venn diagram showing number of proteins detected in vCO and E370K-vCO EVs. C. Volcano plot of protein level of vCO and E370K-vCO EVs, plotting the negative log10 q-values (FDR) of all proteins against their log2 fold change (E370K-vCO EVs vs vCOs). Significantly expressed proteins (q-value < 0.05) are labelled. D. Schematic of acute treatment (12 h) of NPCs with vEVs collected from CTRL and E370K COs (top) and GO terms of upregulated pathways (bottom). E-G. Gene networks of WNT, NOTCH and axon guidance pathways upregulated in NPCs upon treatment with E370K-vEVs

We characterized proteins detected in EVs from CTRL and E370K vCOs (vEVs) (Fig EV6B). Interestingly, E370K-vEVs show a different protein composition with a lower number of different proteins (Fig EV6B and Fig 6B). Since we found LGALS3BP being secreted from ventral COs, we hypothesize that EV-mediated secretion could regulate cell fate decisions in vCOs. (Fig EV6C).

We then performed DE analysis in vEVs. Interestingly we found the secreted molecule SHH (involved in dorso-ventral patterning inducing ventralization(Rallu et al., 2002)) being downregulated in E370K- vEVs suggesting that the low level of SHH carried by EVs in the extracellular space might influence the differentiation of neighboring cells, resulting in a dorsal cell fate. Signaling protein DVL2, known to downregulate the Wnt pathway(Jiang et al., 2015), is downregulated in mutant EVs. The decrease of DVL2 in mutant EVs could lead to an activation of the WNT pathway in neighboring cells, resulting in transcription of dorsal genes(Altmann and Brivanlou, 2001; Chi et al., 2017). EVs secreted by E370K-vCOs also showed dysregulated proteins involved in cell migration and motility (Fig 6E) – PLXNA1, SEMA3G, KIF1A, EPHA2, EPHB2, SLIT1- and proteins of the ECM – HAPLN1, LUM, VCAN –known to modulate cell proliferation and differentiation(Amin and Borrell, 2020; Long et al., 2018; Maeda, 2015). We showed that E370K vCOs have altered the secretion of proteins involved in cellular migration, differentiation, dorso-ventral patterning which could explain the extrinsic effect on these processes. Therefore, we hypothesize that EVs might regulate progenitors’ cell fate and motility, resulting in the dorsal identity observed in ventral E370K neurons.

To this end, we treated NPCs in 2D acutely (12 hours) with EVs isolated from control and E370K vCOs (Fig 6D) and identified upregulated genes in the recipient cells. Interestingly, the RNA-seq analysis performed on NPCs revealed upregulation of WNT and NOTCH pathways (gene networks in Fig 6E and F) after treatment with E370K-vEVs, showing a similar profile with the E370K neural progenitors isolated from vCOs and analyzed by single-cell RNA-Seq (Fig 2F). These results suggest that mutant EVs can change the transcriptomic profile of NPC (Fig EV6D) and activate pathways involved in cell dorso-ventral patterning and cell-cell communication resulting in alteration in their fate(Kumar et al., 2019)(Hayward et al., 2008; Lai, 2004; De Strooper and Annaert, 2001). Moreover, upon treatment with E370K-vEVs, NPCs show upregulation of genes involved in axon guidance (gene network in Fig 6G), as it was found in E370K neurons isolated from vCOs and analyzed by single-cell RNA-Seq (Fig 2H). Altogether, these results propose an extrinsic regulation of neuronal differentiation and migration mediated via EVs.

We showed that vCOs with a variation in the ECM protein LGALS3BP reveal dorsal identity and migratory defect that can lead to NDDs. It has been shown that EVs can also transport ECM remodeling cargoes, like matrix metalloproteinases, that regulate cell growth, differentiation, and cell migration(Nawaz and Fatima, 2017). Altered deposition of collagens mediated via EVs can change the ECM composition and might lead to NDDs(Amin and Borrell, 2020). To better correlate the dysregulated network of proteins altered in E370K-EVs with NDDs, we identified proteins whose genes are associated with cortical malformations (CMs), autism spectrum disorder (ASD), and epilepsy (EP). We identified multiple proteins altered in E370K-EVs compared to control EVs that are mostly associated with microtubule organization, morphogenesis and synaptic activity highlighting the key role of EVs-mediated signaling in NDDs (Fig EV6E-G).

## Discussion

Defects in interneurons specification, migration, and/or recruitment might cause an excitatory/inhibitory imbalance in neuronal circuits, characteristic of neurological disorders, such as epilepsy and ASD. For example, we have recently found that individuals affected with EPM1 (Progressive Myoclonus Epilepsy type1) show loss GABA synaptic terminals(Buzzi et al., 2012) and recruitment of INs is altered in EPM1-hCAs(Di Matteo et al., 2020). Moreover, some ASD patients have reduced GABAergic neurons in the cortex(Ariza et al., 2018; Puts et al., 2016), and organoids derived from patients with ASD show dysregulation in genes involved in GABAergic interneuron differentiation and migration(Wang et al.)(Mariani et al., 2015). Other studies performed in cerebral organoids with mutation of ASD-associated genes revealed alteration in of both GABAergic and excitatory neuron developmental trajectories.(Paulsen et al., 2022; Villa et al., 2022) In this work, we showed that vCOs with a mutation in the *LGALS3BP* gene exhibit alterations in the dorso-ventral patterning resulting in dorsal identity. Previous studies in mice showed that the dorsoventral patterning during brain development is regulated by *Pax6*, *Gli3*, *Shh*(Fuccillo et al., 2006; Theil et al., 1999; Tole et al., 2000). *Gli3* and *Pax6* mutant mice show ventral identity acquired by dorsal pallial tissue expressing ectopic *Dlx2*(Theil et al., 1999; Tole et al., 2000). Mice carrying a deletion in the *Gli3* gene show ectopic ventral telencephalic identity in the dorsal forebrain, probably given by an alteration in the *Wn*t pathway(Tole et al., 2000). We previously described the extracellular function of LGALS3BP in regulating proliferation, NPCs delamination, neuronal distribution, and migration(Kyrousi et al., 2021). In this study, we propose an extrinsic regulation mediated by LGALS3BP in ventral progenitor specification. In the ECM, LGALS3BP binds Galectin-3 (LGALS3), forming a complex that interacts with membrane tetraspanins CD9 and CD82, activating the Wnt/b-catenin signaling(Lee et al., 2010; Pikkarainen et al., 2017). This pathway triggers the gene expression of Wnt target genes in the dorsal forebrain(Chi et al., 2017). One possible scenario is that the E370K variation activates the Wnt/b-catenin signaling in ventral COs. The activation of the signaling might lead to the expression of *PAX6* and repression of *NKX2-1*, resulting in the expression of other cortical dorsal genes (EOMES, TBR1, and SATB2) that define the dorsal identity of E370K vCOs. Moreover, Wnts can act as paracrine molecules, affecting neighboring cells. We showed that the extracellular environment influences molecular identity and neurons specification in vMOs. In vMOs, control and mutant progenitors share similar transcriptomic features; however, after the IPs transition, they reveal their intrinsic program identity (ventral for control and more dorsal for E370K). Final IN identity is refined at later stages in interaction with the local extracellular environment. Indeed, we showed that the proximity of control cells has partially reverted the dorsal identity of E370K cells.

Because of neuronal cell fate changes, the migration of neurons is affected, indeed the dorsal identity acquired by E370K neurons could result in a different response to extrinsic attractive/repulsive stimuli. We described the migratory behavior of E370K cells, finding a significant decrease in velocity that become more similar to radially migrating excitatory neurons as shown previously(Klaus et al.). These cells reflect a more tortuous migratory behavior as it was previously described in migrating neurons derived from an individual with PH (with a mutation in *DCHS1* and *FAT4*)(Klaus et al.). Interestingly, ventrally-derived neurons in E370K show similar molecular signatures with the altered neurons found in DCHS1 and FAT4 patients’ organoids, suggesting shared dysregulated pathways in ectopic neurons accumulating below the white matter in patients with PH. Future studies with cells derived from PH patients with different mutations will be essential to understand if there are common signatures for all PH neurons, providing additional tools to target a specific subpopulation of affected neurons.

We showed that EVs carry secreted molecules that drive dorso-ventral patterning and neuronal migration. Treatment with E370K-vEVs can activate WNT and NOTCH pathways as well as genes associated with neuronal migration and motility in NPCs, confirming their role in cell fate and migration.

## Conclusion

In conclusion, we propose that the EVs can modulate neuronal progenitors fate regulating the dorso- ventral patterning and instruct interneuron migration during brain development. Indeed, a mutation in the ECM component LGALS3BP alters cell fate specification and migration via regulation mediated by factors secreted by EVs. We suggest that the extracellular environment has a key role in the maintenance of excitatory/inhibitory balance during brain development. Thus, the alteration of cell identity and migration caused by the LGALS3BP E370K variation could lead to the disruption of the E/I balance, which might lead to neurodevelopmental and neuropsychiatric disorders.

## Materials and methods

### IPSCs culture

Induced pluripotent stem cells (iPSCs) reprogrammed from NuFF3-RQ human new-born foreskin feeder fibroblasts (GSC-3404, GlobalStem) (Cárdenas et al., 2018) were cultured on Matrigel (Corning) coated plates (Thermo Fisher, Waltham, MA, USA) in mTesR1 basic medium supplemented with 1x mTesR1 supplement (Stem Cell Technologies, Vancouver, Canada) at 37°C, 5% CO2 and ambient oxygen level. Passaging was done using accutase (Stem Cell Technologies) treatment.

### CRISPR genome editing for generation of mutant iPSCs lines

For CRISPR genome editing for the generation of mutant iPSCs, one control iPSC line was used to generate isogenic control and mutant lines as described in Kyrousi et al., 2021.

### Generation of labeled iPSC line

The GFP and RFP-labeled iPSC lines were generated using the piggyBac transposase (1ug) and PB- GFP (1ug)(Di Matteo et al., 2020) and PB-RFP (1ug) nucleofection(Chen and LoTurco, 2012). Single cells of iPSCs were transfected with the Amaxa Nucleofector 2b (program B-016). GFP and RFP colonies were picked and cultured on Matrigel (Corning/VWR International, 354234) coated plates in mTeSR1 basic medium (Stem Cell Technologies, 85850) supplemented with 1× mTeSR1 supplement (Stem Cell Tech- nologies, 85850) at 37°C and 5% CO2.

### Generation of dvCAs and vCOs

Dorso-ventral CAs and vCOs were generated according to Bagley et al, 2017(Bagley Joshua A , Reumann Daniel , Bian Shan, 2017). Embryoid bodies (EBs) were guided to generate ventral and dorsal identities. iPSCs from isogenic control and mutant lines, were dissociating into single cells using Accutase (Sigma-Aldrich, A6964) and approximately 9,000 cells were transferred to one well of an ultra-low-attachment 96-well plate (Corning). Five days later, during the neuronal induction, to induce brain regional patterning, EBs were treated individually with SAG (1:10,000) (Milli- pore, 566660) + IWP-2 (1:2,000) (Sigma-Aldrich, I0536) for ventral identity and with cyclopamine A (1:500) (Calbiochem, 239803) for dorsal identity. After 7 days, one ventral EB and one dorsal EB were embedded together into the same Matrigel (Corning/VWR International, 354234) droplet in order to form a fused organoid. The ventrally patterned EBs were embedded separately, each in one drop of Matrigel. After this point, the generation of vCOs followed methods according to Lancaster & Knoblich, 2017(Lancaster et al., 2017).

### Generation of vMOs

Ventral MOs were generated according to Bagley et al, 2017. iPSCs from isogenic control and from GFP labelled E370k line, were dissociating into single cells using Accutase (Sigma-Aldrich, A6964) and approximately 4,500 cells of each line were mixed together and transferred to one well of an ultra-low-attachment 96-well plate (Corning), in a ratio of 1:1. The protocols continues as described in “Generation of dvCAs and vCOs”.

### Immunohistochemistry

For IHC, sections were post-fixed using 4% PFA for 10 min and permeabilized with 0.3% Triton for 5 min. After post-fixation and permeabilization, sections were blocked with 0.1% Tween, 10% Normal Goat Serum (Biozol, VEC-S-1000). Primary and secondary antibodies were diluted in blocking solution. Nuclei were visualized using 0.5 mg/ml 4,6-diamidino-2-phenylindole (DAPI) (Sigma-Aldrich, D9542). Immunostained sections were analyzed using Leica TCS SP8 Confocal microscope (Leica, Germany). For nuclear antibodies, before the post-fixation step, sections were incubated in a freshly made 10 mM citric buffer (pH 6) in a microwave for 1 min at 720 W and for 10 min at 120 W and then left it to cool down for 20 min at RT. Cells quantifications were performed with the ImageJ software and analyzed with RStudio or GraphPad.

### hCOs 3D immunohistochemistry and tissue clearing

The dvCAs 3D immunohistochemistry and tissue clearing were performed following Masselink et al.,2019. For immunohistochemistry, fused organoids were fixed in 4% PFA overnight at 4C. They were incubated on a shaker for 2 days in PBS-TxDBN solution (10%PBS10X, 2% TX100, 20% DMSO, 5%BSA, 0.05% NaN). Primary and secondary antibodies were diluted in PBS-TxDBN solution and incubated for 4 days. For the following dehydration step, the organoids were transferred sequentially in 30%, 50%, 70% and 2x 99.7% in 1-Propanol(Sigma Cat.W292818):1xPBS (pH:9) solutions. Dehydration is performed at 4C on a gyratory rocker for at least 4h per step. For the refractive index matching, the organoids were transferred into Ethyl-Cinnamate (Sigma Cat. W243000) solution and incubated on a gyratory rocker at RT for at least 1h before recording. The imaging was performed using Leica TCS SP8 Confocal microscope (Leica, Germany). The cells were counted using Imaris software and analyzed with RStudio.

### hCOs 3D time-lapse imaging

For 3D time-lapse imaging, slices of dvCAs were prepared and imaged as described previously(Pilz et al., 2013). Dorso-ventral CAs were sliced at 300-μm thickness on a vibratome (Leica VT1200S) in ice-cold DMEM/F12 (Invitrogen) supplement with sodium bicarbonate, glucose and 10% antibiotics, oxygenated with 100% O2 for 20 min before cutting. The slices were placed on a cell culture insert (Millicell) and further cultured in normal organoid medium. The slices were kept in an atmosphere with 5% CO2 at 37 °C. Live imaging was performed for 48 h using Leica TCS SP8 Confocal microscope (Leica, Germany), taking an image every twenty minutes. The cell movement was tracked using ImageJ software and the Manual Tracking Plugin, and the movement parameters calculated and analyzed in RStudio.

### Single-cell RNA-sequencing library preparation and data analysis

Five 60 days old vCOs and vMCOs were randomly selected from each condition. Single cells were dissociated using StemPro Accutase Cell Dissociation Reagent (Life Technologies), filtered through 30 uM and 20 uM filters (Miltenyi Biotec) and cleaned of debris using a Percoll (Sigma, P1644) gradient. For vMCOs, single cells were FACS sorted to collect control and GFP-labeled E370K single cells. Single cells were resuspended in ice-cold Phosphate-Buffered Saline (PBS) supplemented with 0.04% Bovine Serum Albumin at a concentration of 1000 cells per ul. Single cells were loaded onto a Chromium Next GEM Single Cell 3ʹ chip (Chromium Next GEM Chip G Single Cell Kit, 16 rxns 10XGenomics PN-1000127) with the a Chromium Next GEM Single Cell 3ʹ GEM, Library & Gel Bead Kit v3.1 (Chromium Next GEM Single Cell 3ʹ GEM, Library & Gel Bead Kit v3.1, 4 rxns 10xGenomics PN-1000128) and cDNA libraries were generated with the Single Index Kit T Set A, 96 rxns (10xGenomics PN-1000213) according to the manufacturer’s instructions. Libraries were sequenced using Illumina NovaSeq6000 in 28/8/91bp mode (SP flowcell), quality control and UMI counting were performed by the Max-Planck für molekulare Genetik (Germany). Downstream analysis was performed using the R package Seurat (version 3.2). Cells with more than 2,500 or less than 200 detected genes or with mitochondrial content higher that 10% were excluded as well as genes that were not expressed in at least three cells. We excluded clusters with “glycolysis” identity based on GO terms of cluster-specific markers genes(Bhaduri et al., 2020)(Kanton et al., 2019). Normalization of gene expression was done using a global-scaling normalization method (“LogNormalize”, scale.factor = 10000) and the 2000 most variable genes were selected (selection method, “vst”) and scaled (mean = 0 and variance = 1 for each gene) before principal component analysis. The “FindNeighbors” and “FindClusters” functions were used for clustering with resolution of 0.5 and UMAP for visualization. Clusters were grouped based of the expression of known marker genes and differentially expressed gene identified with the “FindAllMarkers” function. The PCA analysis was used to determine pseudo-differentiation axis of telencephalic cells. For pseudotime analysis, we used Monocle3 algorithm(Trapnell et al., 2014) which identifies the overall trajectory of gene expression changes.

### FACS analysis

Single cells obtained after dissociation of 60 days old vMOs where collected for FACS analysis (“see Single-cell RNA-sequencing library preparation and data analysis”), in order to sort control and GFP- labeled E370K for single-cell RNA-seq analysis. FACS analysis was performed at a FACS Aria (BD) in BD FACS Flow TM medium, with a nozzle diameter of 100 µm. For each run, 10,000 cells were analyzed.

### Dorso-ventral identity model

The model implemented in the bmrm R package was used to classify the identity of cells with genes showing minimal expression (10,407), as described in Oberst et al., 2019l(Oberst et al., 2019). The model was trained with a subset of dorsal and ventral control cells and the 30 most-weighted genes were used for fold cross validation of additional dorsal and ventral control cells and prediction of dorsal and ventral mutant cells. The same method was applied to build a model to classify control and mutant cells, selecting for 100 most-weighted genes.

### EVs collection and analysis

EVs were collected from conditioned media from ventral and dorsal COs by the following steps: centrifugation in 300 g for 15 mins, supernatant centrifugation in 2000g for 10 mins, supernatant centrifugation in 10.000 g for 30 mins, supernatant centrifugation in 100.000 g for 120 mins and pellet wash with 1x PBS and centrifugation in 100.000 g for 60 mins. Alternatively, miRCURY Exosome Cell/Urine/CSF Kit (Qiagen, 76743) was used to isolate EVs from conditioned medium according to the manufacturer instructions.

### Sample preparation for mass spectrometry

Purified EVs were lysed in RIPA buffer (150mM NaCl, 50mM Tris pH8, 0.1% DOC, 0.1% SDS, 0.1% NP40). 10 ug of protein for each sample was subjected to the modified FASP protocol (Wiśniewski et al., 2009). Briefly, the protein extract was loaded onto the centrifugal filter CO10 kDa (Merck Millipore, Darmstadt, Germany), and detergent were removed by washing five times with 8M Urea (Merck, Darmstadt, Germany) 50mM Tris (Sigma-Aldrich, USA) buffer. Proteins were reduced by adding 5mM dithiothreitol (DTT) (Bio-Rad, Canada) at 37degrees C for 1 hour in the dark. To remove the excess of DTT, the protein sample was washed three times with 8M Urea, 50mM Tris. Subsequently protein thiol groups were blocked with 10mM iodoacetamide (Sigma-Aldrich, USA) at RT for 45 min. Before proceeding with the enzymatic digestion, urea was removed by washing the protein suspension three times with 50mM ammonium bicarbonate (Sigma-Aldrich, Spain). Proteins were digested first by Lys- C (Promega, USA) at RT for 2 hours, then by trypsin (Premium Grade, MS Approved, SERVA, Heidelberg, Germany) at RT, overnight, both enzymes were added at an enzyme-protein ratio of 1:50 (w/w). Peptides were recovered by centrifugation followed by two additional washes with 50mM ammonium bicarbonate and 0.5M NaCl (Sigma-Aldrich, Switzerland). The two filtrates were combined, the recovered peptides were lyophilized under vacuum. Dried tryptic peptides were desalted using C18-tips (Thermo Scientific, Pierce, USA), following the manufacture instructions. Briefly, the peptides dissolved in 0.1%(v/v) formic acid (Thermo scientific, USA) were loaded onto the C18-tip and washed 10 times with 0.1 % (v/v) formic acid, subsequently the peptides were eluted by 95% (v/v) acetonitrile (Merck, Darmstadt, Germany), 0.1% (v/v) formic acid. The desalted peptides were lyophilized under vacuum. The purified peptides were reconstituted in 0.1% (v/v) formic acid for LC-MS/MS analysis.

### MS data acquisition

Desalted peptides were loaded onto a 25 cm, 75 µm ID C18 column with integrated nanospray emitter (Odyssey/Aurora, ionopticks, Melbourne) via the autosampler of the Thermo Easy-nLC 1000 (Thermo Fisher Scientific) at 60 °C. Eluting peptides were directly sprayed onto the timsTOF Pro (Bruker Daltonics). Peptides were loaded in buffer A (0.1% (v/v) formic acid) at 400 nl/min and percentage of buffer B (80% acetonitril, 0.1% formic acid) was ramped from 5% to 25% over 90 minutes followed by a ramp to 35% over 30 minutes then 58% over the next 5 minutes, 95% over the next 5 minutes and maintained at 95% for another 5 minutes. Data acquisition on the timsTOF Pro was performed using timsControl. The mass spectrometer was operated in data-dependent PASEF mode with one survey TIMS-MS and ten PASEF MS/MS scans per acquisition cycle. Analysis was performed in a mass scan range from 100-1700 m/z and an ion mobility range from 1/K0 = 0.85 Vs cm-2 to 1.30 Vs cm-2 using equal ion accumulation and ramp time in the dual TIMS analyzer of 100 ms each at a spectra rate of 9.43 Hz. Suitable precursor ions for MS/MS analysis were isolated in a window of 2 Th for m/z < 700 and 3 Th for m/z > 700 by rapidly switching the quadrupole position in sync with the elution of precursors from the TIMS device. The collision energy was lowered as a function of ion mobility, starting from 45 eV for 1/K0 = 1.3 Vs cm-2 to 27eV for 0.85 Vs cm-2. Collision energies were interpolated linear between these two 1/K0 values and kept constant above or below these base points. Singly charged precursor ions were excluded with a polygon filter mask and further m/z and ion mobility information was used for ‘dynamic exclusion’ to avoid re-sequencing of precursors that reached a ‘target value’ of 14500 a.u. The ion mobility dimension was calibrated linearly using three ions from the Agilent ESI LC/MS tuning mix (m/z, 1/K0: 622.0289, 0.9848 Vs cm-2; 922.0097 Vs cm-2, 1.1895 Vs cm-2; 1221.9906 Vs cm-2, 1.3820 Vs cm-2).

### Raw data analysis of MS measurements

Raw data were processed using the MaxQuant computational platform (version 1.6.17.0)(Tyanova et al., 2016) with standard settings applied for ion mobility data(Prianichnikov et al., 2020). Shortly, the peak list was searched against the Uniprot database of Human database (75069 entries, downloaded in July 2020) with an allowed precursor mass deviation of 10 ppm and an allowed fragment mass deviation of 20 ppm. MaxQuant by default enables individual peptide mass tolerances, which was used in the search. Cysteine carbamidomethylation was set as static modification, and methionine oxidation, deamidation and N-terminal acetylation as variable modifications. The match-between-run option was enabled, and proteins were quantified across samples using the label-free quantification algorithm in MaxQuant generating label-free quantification (LFQ) intensities. The mass spectrometry proteomics data have been deposited to the ProteomeXchange Consortium via the PRIDE(Perez-Riverol et al., 2019).

### Bioinformatic analysis

For the proteomic characterization in EVs, 3427 proteins were quantified. Proteins that were consistently detected in 2 of the 3 technical replicates per each condition were retained. Downstream analysis was performed using R. The LFQs values were log2-transformed. Missing values were imputed using the R package DEP (version 1.15.0) and replaced by random values of a left-shifted Gaussian distribution (shift of 1.8 units of the standard deviation and a width of 0.3). Differentially expression (DE) analysis was performed on the imputed data using Student’s t-Test. Proteins with log2 fold change values (log2FC) ≥ 1 and ≤ -1 and with an FDR-corrected q-value < 0.05 were considered as differentially expressed. The gene-disease associations analysis was performed using DisGeNET (https://www.disgenet.org/search Piñero et al., 2017).

### Bulk-RNA-sequencing

RNA-seq was performed on 10ng of total RNA collected from 3 independent wells of NPCs from a 24well plate. NPCs were not treated with EVS or treated for 12h with EVs collected by ultracentrifugation from 25 ml of conditioned medium collected from 28 to 37 days in culture COs (control ventral, EPM1 ventral, control dorsal and EPM1 dorsal COs). NPCs were lysed in 1ml Trizol(Qiagen)/well and RNA was isolated employing RNA Clean & Concentrator kit (Zymo Research) including digestion of remaining genomic DNA according to producer’s guidelines. RNA was further processed according to(Cernilogar et al., 2019). Briefly, cDNA synthesis was performed with SMART-Seq v4 Ultra Low Input RNA Kit (Clontech cat. 634888) according to the manufacturer’s instruction. cDNA was fragmented to an average size of 200–500 bp in a Covaris S220 device (5 min; 4°C; PP 175; DF 10; CB 200). Fragmented cDNA was used as input for library preparation with MicroPlex Library Preparation Kit v2 (Diagenode, cat. C05010012) and processed according to the manufacturer’s instruction. Libraries were quality controlled by Qubit and Agilent DNA Bioanalyzer analysis. Deep sequencing was performed on a HiSeq 1500 system according to the standard Illumina protocol for 50 bp paired-end reads with v3 sequencing reagents.

### RNAseq analysis

Paired end reads were aligned to the human genome version GRCh38 using STAR v2.6.1d(Dobin et al., 2013) with default options “--runThreadN 32 --quantMode TranscriptomeSAM GeneCounts -- outSAMtype BAM SortedByCoordinate”. Reads-per-gene counts were imported in R v4.1.0. Bioconductor package DESeq2 v1.32.0(Love et al., 2014) was used for differential expression analysis. Only genes with read counts>1 were considered. Significantly changed genes were determined through pairwise comparisons using the DESeq2 results function (log2 fold change threshold=1, adjusted p-value <0.05). Heatmaps with differentially expressed genes were plotted with pheatmap v1.0.12 and RColorBrewer v1.1-2 using rlog-normalized expression values.

### Enrichment analysis

GO term analysis of differentially-expressed genes in mutant patterned hCOs was tested using the FUMA algorithm(Watanabe et al., 2017) by inserting the DE gene lists from all the cell populations into the GENE2FUNC software (FDR<0.05) or STRING.

**Table 1.** Immunostaining antibodies

| Primary antibodies |  |  |  |  |  |
| --- | --- | --- | --- | --- | --- |
| Antibody | Host | Company | Catalogue No. | Lot No. | Dilution |
| NKX2.1 | Mouse IgG1 | Merk<br>Millipore | MAB5460 | 3160139 | 1:500 |
| TBR1 | Rabbit | Abcam | Ab31940 | GR3217067-1 | 1:500 |
| SOX2 | Rabbit | Cell Signaling | 27485 | 2 | 1:500 |
| LGALS3BP | Mouse IgG1 | eBioscience | BMS146 | - | 1:100 |
| PAX6 | Rabbit | Biolegend | PRB-278p | B244513 | 1:500 |
| MEIS2 | Mouse IgG1 | Santa Cruz<br>Biotechnology | SC 101850 | H0817 | 1:300 |
| EOMES | Rabbit | Abcam | ab23345 | GR33045451 | 1:500 |
| SATB2 | Mouse IgG1 | Abcam | Ab51502 | GR2075794 | 1:500 |
| CALB2 | Rabbit | Swang | 7697 | - | 1:1000 |
| GFP | Chicken | Aves Lab | GFP-1020 | 697986 | 1:1000 |

| Secondary antibodies |  |  |  |  |  |
| --- | --- | --- | --- | --- | --- |
| Antibody | Host | Company | Catalogue No. | Lot No. | Dilution |
| AlexaFluor647 | Rabbit | ThermoFisher<br>Scientific | A21244 | 2086730 | 1:1000 |
| AlexaFluor488 | Chicken | ThermoFisher<br>Scientific | A11039 | 2147635 | 1:1000 |
| AlexaFluor546 | Mouse IgG1 | ThermoFisher<br>Scientific | A21123 | 1722393 | 1:1000 |
| AlexaFluor488 | Rabbit | ThermoFisher<br>Scientific | A11008 | 2079383 | 1:1000 |

## Acknowledgments

we thank Maik Ködel, Anthi Krontira, Pablo Lopez, Cristiana Cruceanu, Filippo Cernilogar, Sylvain Moser, Francesco Di Matteo, Andrea Forero, Veronica Pravata, Giovanna Berto for technical help and critical discussion.

## Author contributions

Conceptualization: SC, FP. Methodology: FP, NB, CK, RDG, RB. Investigation: SC, FP, NB, RDG. Visualization: FP. Funding acquisition: SC. Supervision: SC, DJ. Writing: original draft: FP; review & editing: SC, FP.

## Expanded View Figure legends

**Fig EV1.**
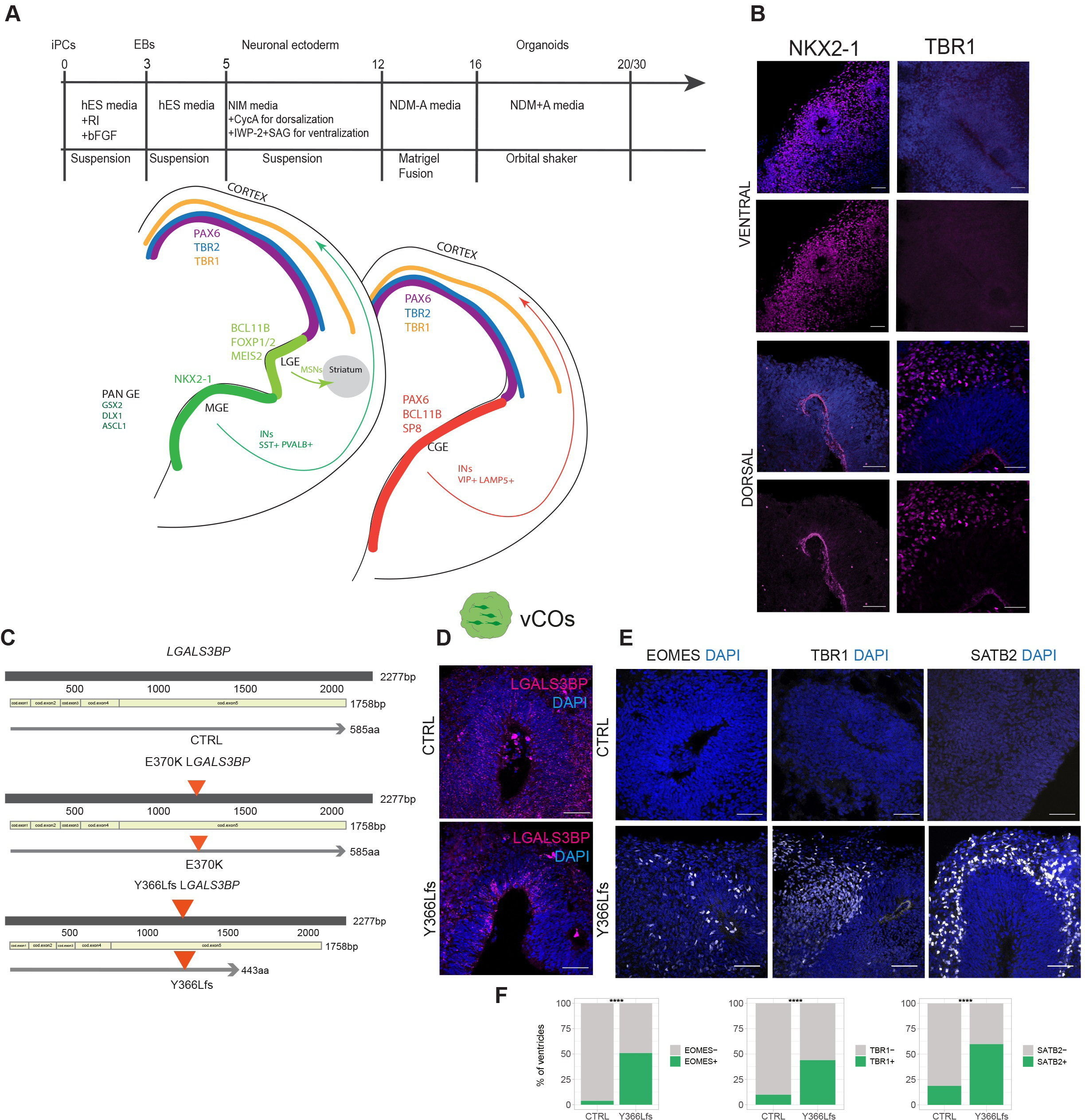
A. Schematic of differentiation protocol used for vCOs and dvCAs (top) and Schematic of tangential migration of interneurons from MGE and CGE to the cortex and markers used for forebrain regional characterization. B. Representative immunostaining for regional markers in dvCAs (NKX2.1 for ventral and TBR1 for dorsal regions). Scale bar: 50 µm. C. Schematic representation of the E370K and Y366Lfs mutant iPSC lines generated using the CRISPR/Cas9 genome editing in control iPSCs. D. Representative immunostaining of CTRL and Y366Lfs vCOs for SOX2 (green) and LGALS3BP (magenta). Scale bar: 50 µm. E. Representative immunostaining of CTRL and Y366Lfs vCOs for cortical markers of IPs (EOMES) of deep layer neurons (TBR1) and upper layer neurons (SATB2). Scale bar: 50 µm. F. Quantification of TBR1+cells in ventral region of CTRLv-CTRLd and E370v-CTRLd dvCAs. Box plots show median and interquartile range. Statistical significance was based on Mann-Withney U test*p<0.05, ***p<0.001. n of organoids: CTRLv-CTRLd =5, LGALS3BPv-CTRLd=9, from at least 2 different batches.

**Fig EV2.**
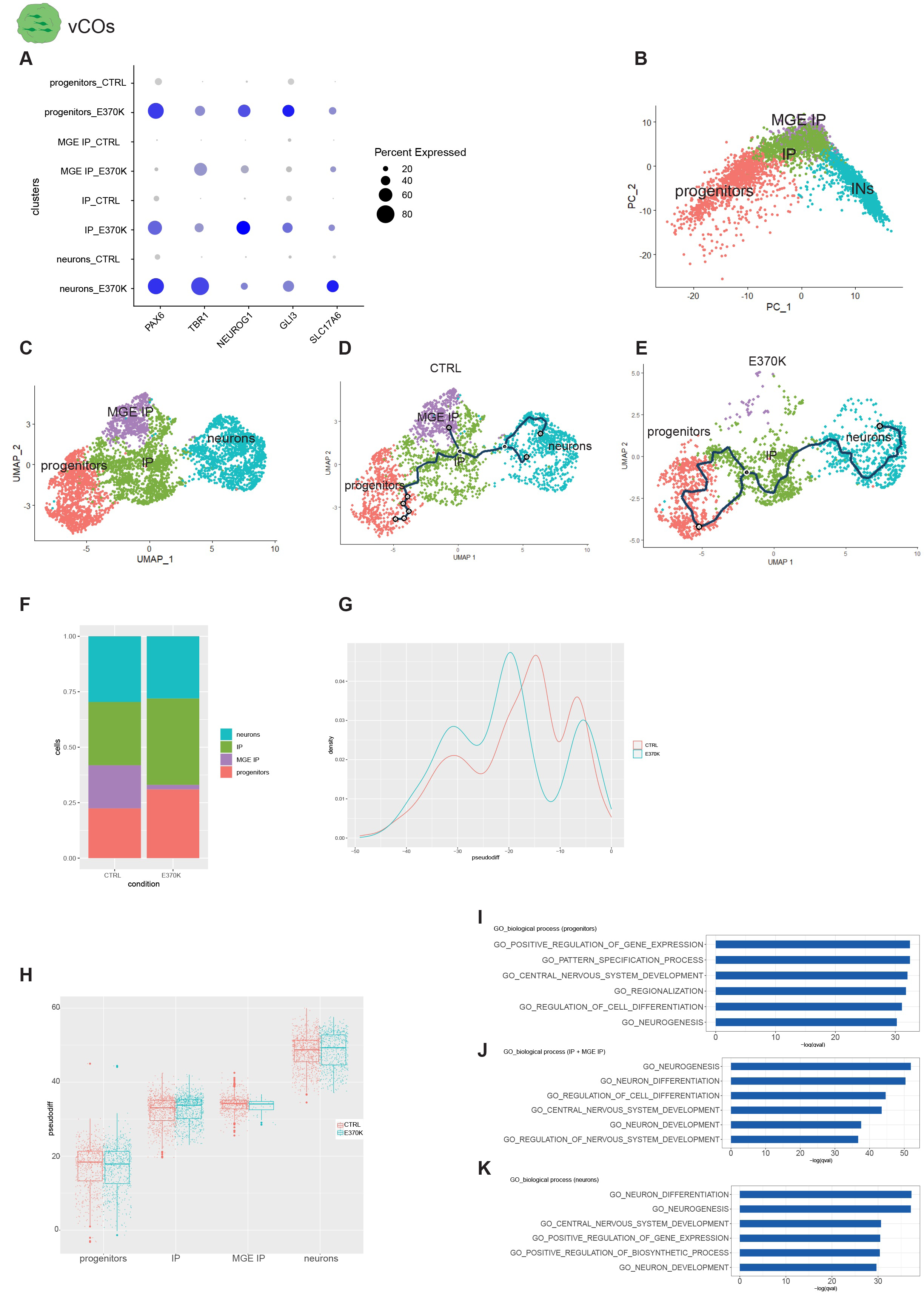
A. Dotplot of cortical genes expressed in vCO clusters. B. PCA visualization and of CTRL and E370K vCOs, showing the pseudo-differentiation axis from progenitors to INs. C. UMAP visualization of scRNA-seq ventral telencephalic cells in CTRL and E370K vCOs. D-E. UMAP visualization of pseudo-differentiation trajectories in CTRL vCOs and in E370K clusters from progenitors to INs. F. Bar plot showing cell proportion in CTRL and E370K vCOs. G. Density plot showing cell distribution along pseudo-differentiation axis in CTRL and E370K vCOs. H. Box plot showing cell distribution along pseudo-differentiation axis in CTRL and E370K vCOs per each cluster. I-K. GO enrichment for DE genes in progenitors, IP and MGE and INs. GO for biological process are reported.

**Fig EV3.**
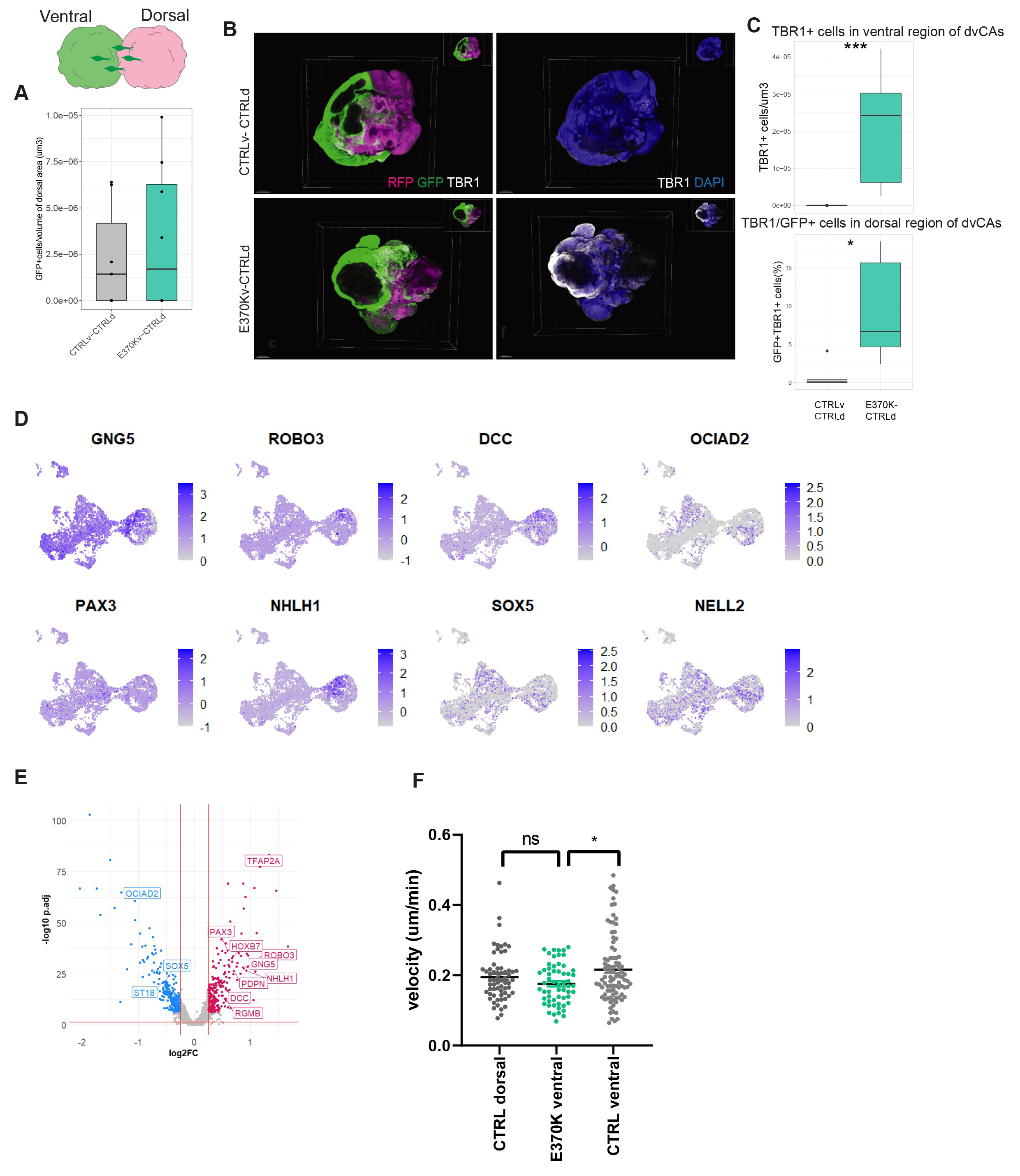
A. Quantification of the number of GFP+ventral cells migrated from ventral to dorsal in dvCAs. Box plots show median and interquartile range. Significance was based on the Mann-Withney U test. Every dot in the plots refers to analyzed vCAs generated in at least 2 independent batches. B. Representative 3D immunostaining of CTRLv-CTRLd and E370Kv-CTRLd dvCAs for GFP and TBR1. Scale bar: 500 µm. C. Quantification of TBR1+cells in ventral region of CTRLv-CTRLd and E370v-CTRLd dvCAs. Box plots show median and interquartile range. Statistical significance was based on Mann-Withney U test*p<0.05, ***p<0.001. n of organoids: CTRLv-CTRLd =5, LGALS3BPv-CTRLd=9, from at least 2 different batches (top) and quantification of migrated GFP+ventral cells expressing TBR1 in dorsal region of CTRLv-CTRLd and E370v-CTRLd dvCAs. Box plots show median and interquartile range. Statistical significance was based on Mann-Withney U test*p<0.05, ***p<0.001. n of organoids: CTRLv-CTRLd =5, LGALS3BPv-CTRLd=9, from at least 2 different batches (bottom). D. Feature plot depicting the expression of genes associated with PH (GNG5, ROBO3, DCC, OCIAD2, PAX6, NHLH1, SOX5, NELL2) in vCOs. E. Volcano plot showing the fold change (CTRL vs E370K) of gene expression of PH-associated genes in E370K neurons. F. Graph showing the comparison between velocity (um/min) of CTRL dorsal neurons (analyzed in Klaus et al., 2019(Klaus et al.)), E370K ventral neurons and CTRL ventral neurons migrating within the dorsal region. Data are shown mean ± SEM Statistical significance was based on the Mann-Withney U test *p<0.05. Every dot in the plots refers to single cells per vCO.

**Fig EV4.**
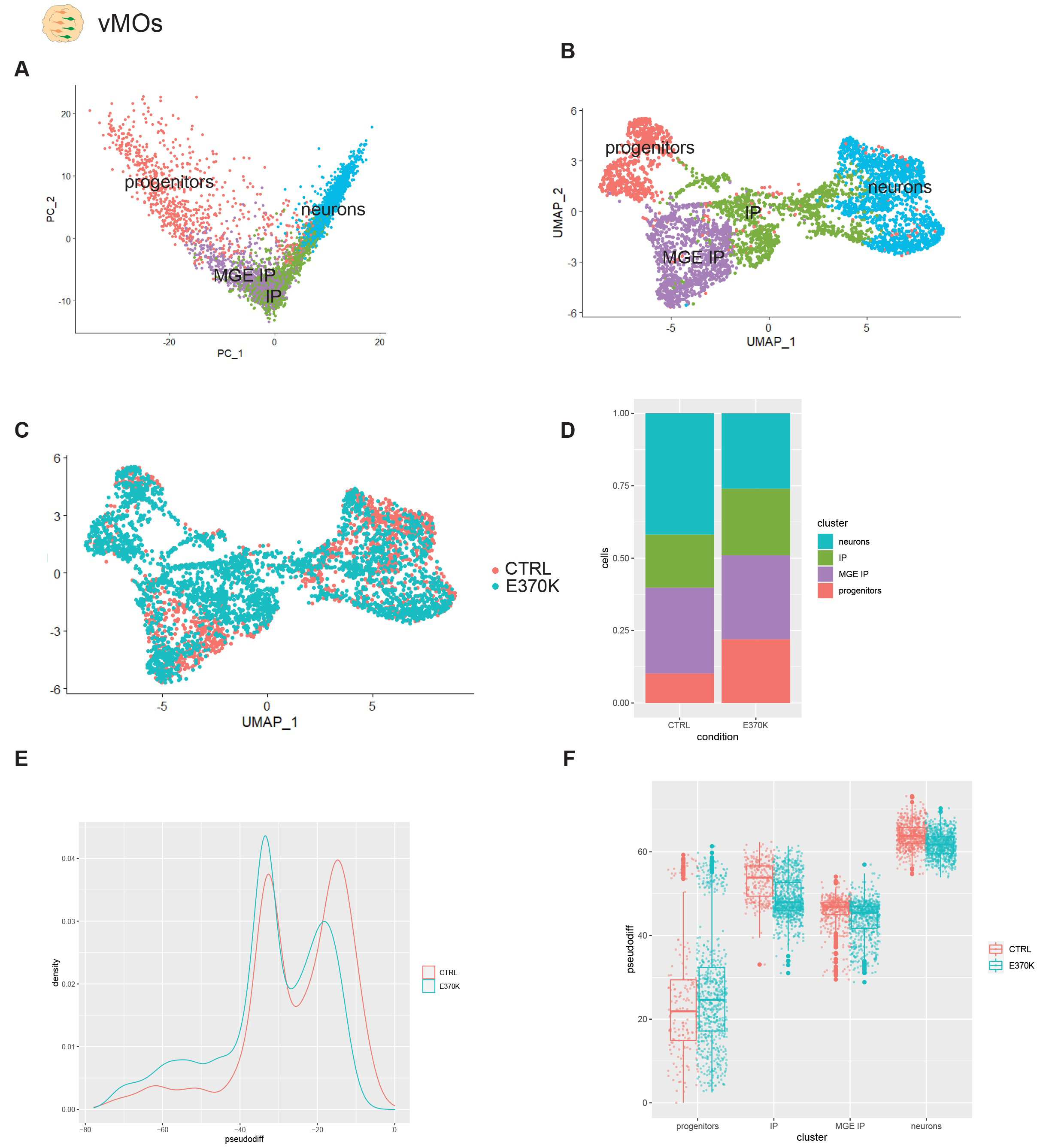
A. PCA visualization and of vMOs, showing the pseudo-differentiation trajectory from progenitors to INs. B-C. UMAP visualization of scRNA-seq ventral telencephalic cells clusters in vMOs. D. Bar plot showing cell proportion in vMOs. E. Density plot showing cell distribution along pseudo-differentiation axis in vMOs. F. Box plot showing cell distribution along pseudo-differentiation axis in vMCOs per each cluster. G. Volcano plot showing the fold change (CTRL vs E370K) of gene expression in progenitors in vMCOs.

**Fig EV5.**
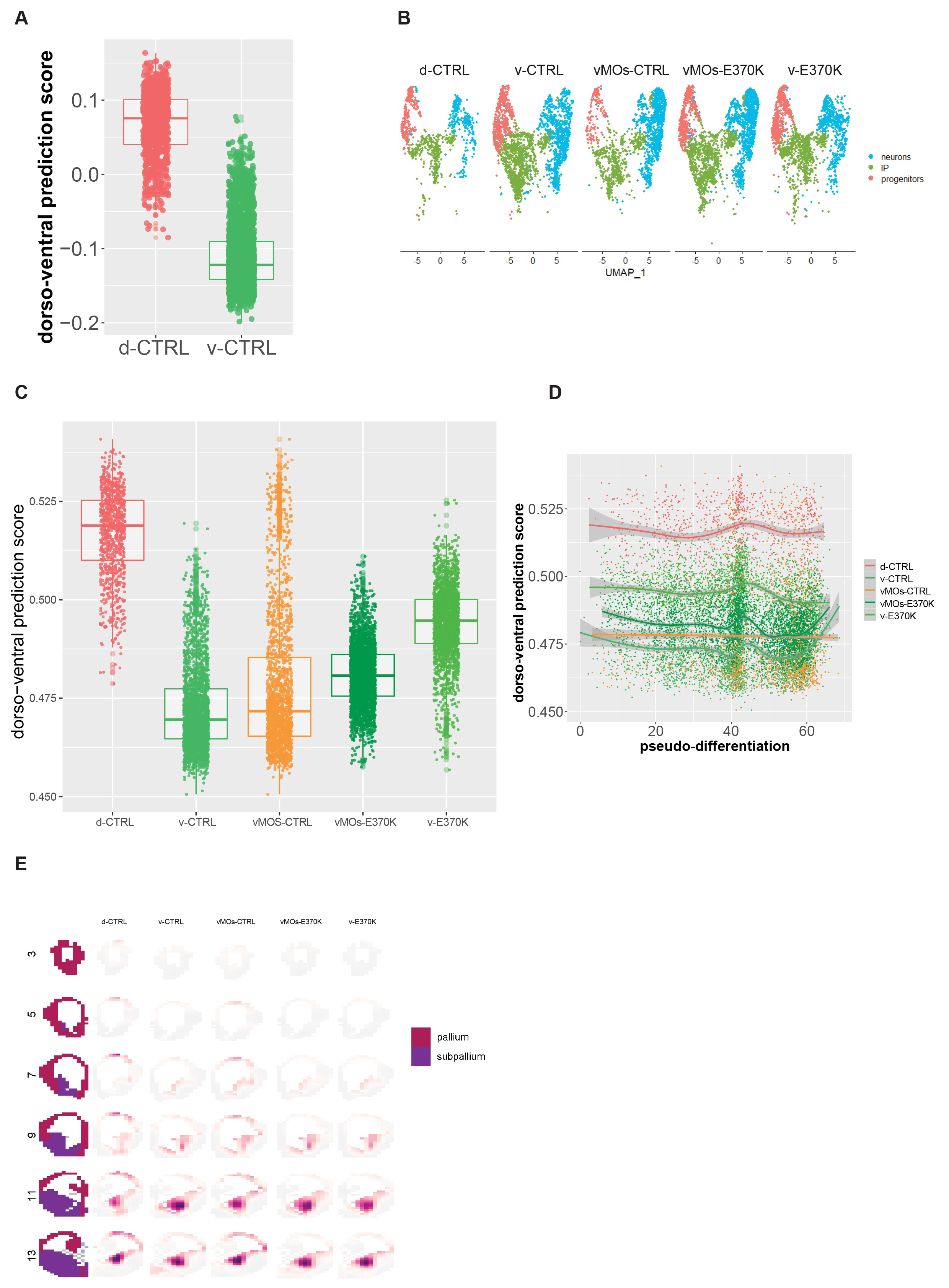
A. Box plot showing the dorso-ventral prediction score based on fold cross validation. B. UMAP visualization of scRNA-seq clusters of CTRL dCOs, CTRL vCOs, E370K vCOs and vMOs. C. Boxplot showing the dorso-ventral prediction score of CTRL dCOs, CTRL vCOs, E370K vCOs and vMOs cells. Box plots show median and interquartile range. D. Density plot showing the distribution of dorso-ventral prediction score of each condition along the pseudo-differentiation axis. E. VoxHunt spatial brain mapping of the scRNA-seq from all conditions onto data from E13.5 mouse brains from the Allen Brain Institute.

**Fig EV6.**
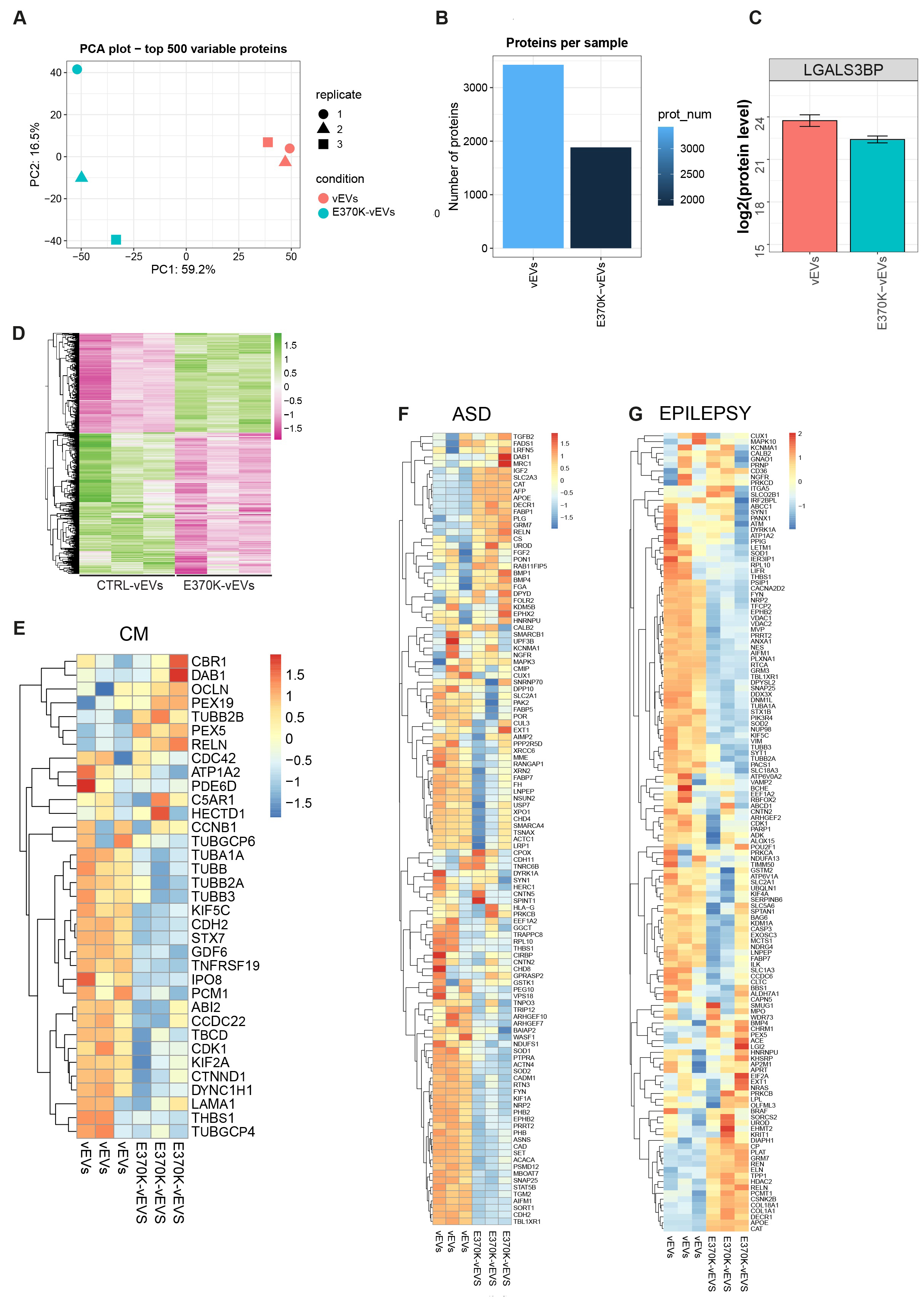
A. PCA plot of protein samples from EVs collected from CTRL and E370K vCOs, based on LFQ intensity of quantified proteins. All the replicates are represented. B. Bar plot showing the number of proteins detected in vEVs and E370K-vEVs. C. Bar plot of LGALS3BP protein level detected in vEVs and E370K-vEVs. Heatmap showing transcriptomic profile changes in NPCs after treatment with E370K-vEVs E-G. Heatmap of DE protein in and E370K-vEVs associated with MCD, epilepsy and ASD.

